# RNA G-quadruplexes mediate cooperativity in HNRNPH binding and splicing regulation

**DOI:** 10.64898/2026.03.03.709289

**Authors:** Kerstin Tretow, Mario Keller, Mikhail Mesitov, Miona Ćorović, Mirko Brüggemann, Jonas Busam, Farica Zhuang, Nadine Körtel, Nicolas Melchior, Anke Busch, Simon Braun, Nadja Hellmann, Heike Hänel, Pol Besenius, Susanne Strand, Yoseph Barash, Stefan Legewie, Friederike Schmid, Julian König, Kathi Zarnack

**Affiliations:** Institute of Molecular Biology (IMB), Ackermannweg 4, 55128 Mainz, Germany; Buchmann Institute for Molecular Life Sciences (BMLS) & Institute of Molecular Biosciences, Goethe University Frankfurt, Max-von-Laue-Str. 15, 60438 Frankfurt, Germany; Theodor Boveri Institute, Julius Maximilians University Würzburg, Biocenter, Am Hubland, 97074 Würzburg, Germany; Department of Computer and Information Science, University of Pennsylvania, 3330 Walnut Street, Philadelphia, PA 19104, USA; Department of Chemistry, Biochemistry, Johannes Gutenberg University Mainz, Hanns-Dieter-Hüsch-Weg 17, 55128 Mainz, Germany; Department of Chemistry, Johannes Gutenberg University Mainz, Duesbergweg 10-14, 55128 Mainz, Germany; Department of Medicine I, University Medical Center of the Johannes Gutenberg University Mainz, Obere Zahlbacher Straße 63, 55131 Mainz, Germany; Department of Genetics, Perelman School of Medicine, University of Pennsylvania, 415 Curie Boulevard, Philadelphia, PA 19104, USA; University of Stuttgart, Institute of Biomedical Genetics (IBMG) & Stuttgart Research Centre Systems Biology (SRCSB), Allmandring 31, 70569 Stuttgart, Germany; Institute of Physics, Johannes Gutenberg University Mainz, Staudingerweg 7-9, 55128 Mainz, Germany

**Keywords:** HNRNPH, alternative splicing, cooperative regulation, RNA structure, RNA G-quadruplex, G4, cancer, iCLIP, RTstop profiling, antisense oligonucleotides, mathematical modelling

## Abstract

Alternative splicing, regulated by RNA-binding proteins (RBPs), enables the generation of diverse transcript isoforms critical for cellular function. However, how RNA secondary structure impacts RBP binding and function remains poorly understood. Here, we unravel how RNA G-quadruplexes (rG4s) facilitate cooperativity in splicing regulation by the RBP heterogeneous nuclear ribonucleoprotein H (HNRNPH). Through high-throughput *in vivo* and *in vitro* studies combined with theoretical modeling, we dissect how rG4s mediate cooperative HNRNPH binding to RNA, ultimately modulating the splicing of hundreds of exons. rG4 unfolding by HNRNPH exposes multiple G-rich binding sites, thereby establishing indirect cooperativity, which is further amplified to achieve switch-like splicing regulation. HNRNPH-mediated regulation is evident in breast cancer patients, with tumors showing rG4-disrupting variants and global HNRNPH alterations, driving distinct splicing patterns that distinguish tumor subtypes. Overall, our findings offer valuable insights into the mechanistic role of RNA secondary structures in cooperative RBP binding and splicing regulation and highlight the clinical relevance of HNRNPH-dependent splicing in cancer.

**Highlights:**

- Hundreds of cassette exons are cooperatively regulated by HNRNPH.
- Unfolding of RNA G-quadruplexes (rG4s) at HNRNPH binding sites facilitates indirect cooperativity in RNA binding.
- Multi-step splicing amplifies the response into highly switch-like regulation.
- rG4-disrupting variants and changing *HNRNPH* expression are associated with breast cancer phenotypes.

## Introduction

RNA regulation is central to gene expression, adding multiple layers of control to when and how genes produce functional proteins. A key process in RNA regulation is splicing, which removes introns and joins exons of precursor messenger RNA (pre-mRNA) and, through alternative splicing, allows to generate distinct mRNA isoforms from a single gene. RNA-binding proteins (RBPs) are crucial regulators of splicing, guiding the splicing machinery to target exons and introns. Yet, while many RBPs are known to influence splicing, the mechanisms underlying their regulation—particularly through RNA secondary structures—remain largely unclear.

RBP binding to RNA often occurs in a cooperative manner whereby the binding of one RBP molecule influences the binding of subsequent molecules, either positively or negatively (Corley, Burns, and Yeo 2020). By contrast, in non-cooperative binding, the RBP’s affinity for its binding sites remains constant, regardless of whether other sites are occupied. Importantly, the cooperativity in RBP–RNA interactions allows for switch-like regulation, facilitating sharp responses to changes in cellular conditions. Through cooperative binding, RBPs can amplify their regulatory effects in RNA splicing and other processes. For example, cooperative binding of RBPs and microRNAs to stem loops in the 3’ untranslated regions (3’ UTRs) of target mRNAs influences their posttranscriptional control (HafezQorani et al. 2016), and RBP cooperativity has been linked to the nuclear export of viral RNAs (Daugherty, Liu, and Frankel 2010). Moreover, the formation of cooperative higher-order RBP complexes on pre-mRNAs has been identified as a critical determinant for individual splicing decisions (Cartegni et al. 1996; Gueroussov et al. 2017). Such cooperative effects are critical for the dynamic control of gene expression, yet their molecular basis remains poorly explored.

Cooperativity in RBP binding can be achieved through direct or indirect mechanisms. In direct cooperativity, two or more RBPs bind to RNA in close proximity and stabilize each other’s binding through physical interactions (Campbell et al. 2012; Cartegni et al. 1996; Gueroussov et al. 2017). Indirect cooperativity can arise through RNA secondary structures, for instance when RBP binding sites are buried within folded RNA. When an RBP unfolds the RNA structure, additional sites may become accessible, leading to a cooperative effect (Becker et al. 2019; Lin and Bundschuh 2015). Although these mechanisms have been described for individual splicing events (Becker et al. 2019; Gueroussov et al. 2017; Lin and Bundschuh 2015; Schaub, Lopez, and Caputi 2007), a comprehensive understanding of cooperativity on a transcriptome-wide scale has yet to be established.

Heterogeneous nuclear ribonucleoprotein H (HNRNPH), referring collectively to the close paralogs HNRNPH1 and HNRNPH2 due to their 96% sequence similarity (Schaub et al. 2007), is an RBP that binds guanine (G)-rich RNA sequences and regulates various aspects of RNA biology (Brownmiller and Caplen 2023). In alternative splicing, HNRNPH has been shown to either promote or repress exon inclusion, depending on its position within the transcript (Braun et al. 2018; Chen and Helfman 1999; Chou et al. 1999; Lefave et al. 2011; Neckles et al. 2019; Wang, Dimova, and Cambi 2007; Yamazaki et al. 2018). We could recently show that HNRNPH represses the alternative splicing of *MST1R* exon 11 in a cooperative manner (Braun et al. 2018).

Prior studies suggested that HNRNPH binding is sensitive to RNA structural features. HNRNPH binds with its three quasi RNA recognition motif (qRRM) domains to G-rich RNA sequences with GGG triplets (Caputi and Zahler 2001; Dominguez et al. 2010). Such G-runs can fold ino RNA G-quadruplexes (rG4s), stable secondary structures formed by stacked G-quartets consisting of four G residues in a planar, cyclic arrangement, stabilized by Hoogsten base-pairing (consensus sequence G_3-5_N_1-7_G_3–5_N_1-7_G_3-5_N_1-7_G_3-5_N_1-7_ where N is any nucleotide) (Dumas et al. 2021). Features like loop length, number of G-quartets and monovalent cations determine rG4 stability (Guiset Miserachs et al. 2016; Lane et al. 2008; Wong and Wu 2003), creating a compact structure that can affect RBP accessibility. Indeed, HNRNPH has been linked to rG4s in different contexts, including its sequestration to rG4s formed by *C9ORF72* G_4_C_2_ repeat expansion (C, cytosine)(Conlon et al. 2016). rG4s have also been proposed to contribute to HNRNPH-mediated splicing regulation (Conlon et al. 2016; Dardenne et al. 2014; Vo et al. 2022), although their impact and mode of action remained unclear. Moreover, a biochemical *in vitro* study resolved the binding of the closely related protein HNRNPF to an rG4 structure in the *CD44* mRNA (Huang et al. 2017). In contrast, other studies suggested that, in combination with RNA helicases, both HNRNPH and HNRNPF prefer binding to unfolded G-runs ((Herviou et al. 2020a; Huang et al. 2017).

Here, we reveal an intricate splicing-regulatory mechanism mediated by cooperative HNRNPH binding and modulated by rG4 structures at hundreds of exons in the human transcriptome. Using high-throughput *in vivo* and *in vitro* experiments combined with theoretical approaches, we demonstrate that HNRNPH binding sites fold into rG4s, which enhance HNRNPH binding and enable cooperative interactions. We propose that HNRNPH recognizes and unfolds the rG4 structures, facilitating indirect cooperativity by exposing multiple G-rich binding sites. This cooperativity is amplified into steep splicing responses at regulated exons, creating switch-like regulation that sensitively follows HNRNPH levels in cells. We demonstrate that rG4-disrupting variants occur in breast cancer patients and that *HNRNPH* levels vary dynamically across tumor subtypes and influence cancer cell proliferation.

## Results

### Hundreds of exons are cooperatively regulated by HNRNPH

We previously found that the alternative splicing of *MST1R* exon 11 is regulated by HNRNPH in a switch-like manner (Braun et al. 2018). To test if this is a transcriptome-wide phenomenon, we performed RNA sequencing (RNA-seq) on a titration series including seven consecutive *HNRNPH* knockdown (KD) (Braun et al. 2018) and three *HNRNPH* overexpression (OE) steps in MCF7 cells. Total HNRNPH protein levels (sum of HNRNPH1 and HNRNPH2) ranged from 46% to 176% of mock-treated control (Ctrl) HNRNPH protein (12 conditions, in triplicates, 54–70 million reads per sample; **Figure 1A, B, Figure S1A; Supplementary Table S1**). *HNRNPH1* mRNA levels in the RNA-seq data showed a strong correlation with HNRNPH protein levels from Western blots (**Figure S1B**). In the KD samples, we also observed a concomitant downregulation of *HNRNPH2*, while housekeeping genes remained unchanged (**Figure S1C**).

**Figure 1.**
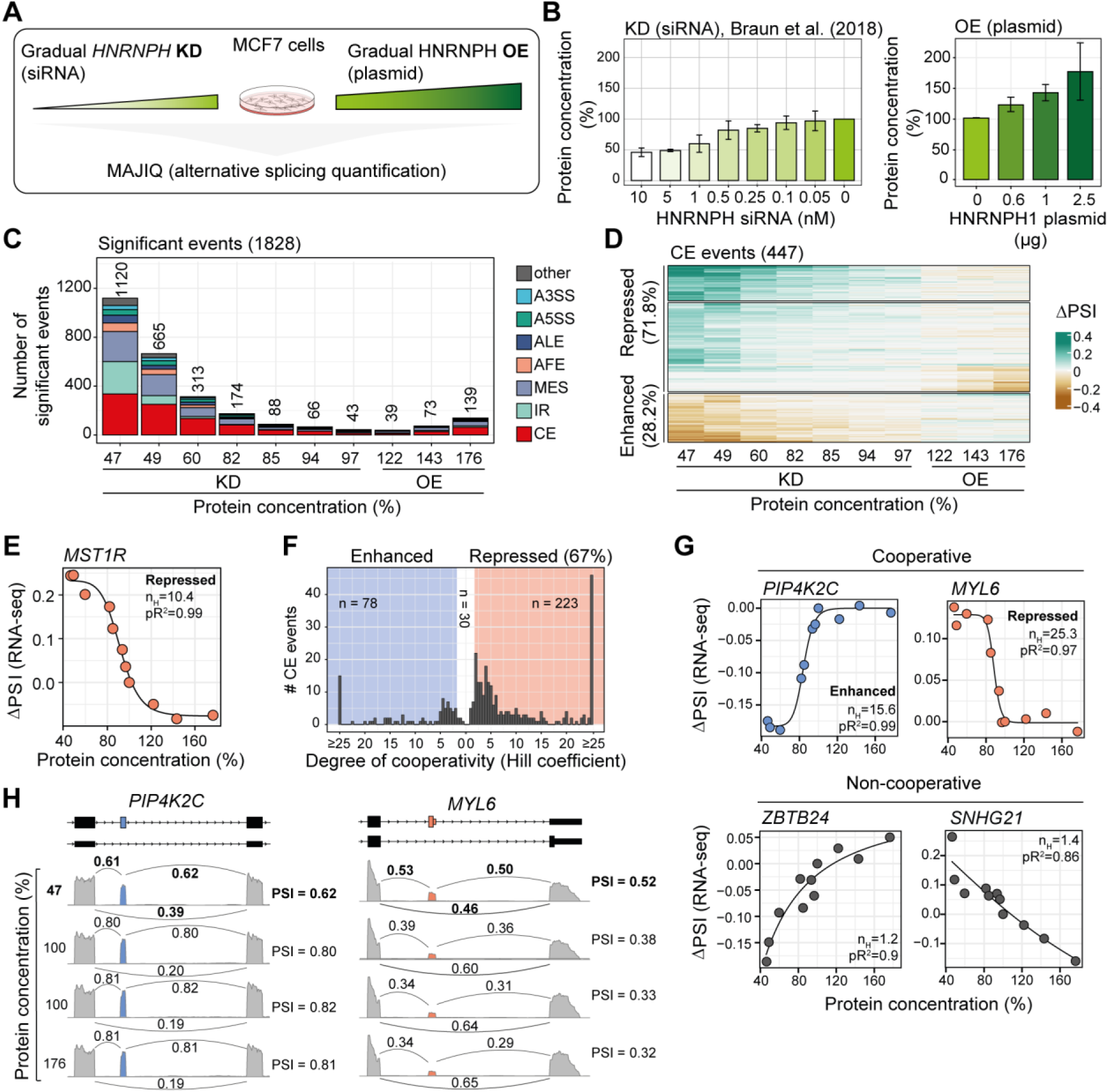
Titration of HNRNPH reveals cooperative regulation of alternative splicing. **(A)** Experimental design of HNRNPH titration experiment: HNRNPH levels were gradually reduced (siRNA) and increased (overexpression constructs). RNA-seq was used to measure expression and splicing changes, Western Blot measured protein levels, and alternative splicing changes were quantified with MAJIQ (Vaquero-Garcia et al. 2016). **(B)** Box plots show the quantification of HNRNPH, measuring knockdown ((Braun et al. 2018); left) and overexpression (right) on the protein level. **(C)** Stacked bar chart shows the number of significantly regulated events per event class for each knockdown and overexpression condition. Event classes include cassette exon (CE), intron retention (IR), multi-exon skipping (MES), alternative last exon (ALE), alternative first exon (AFE), alternative 5’ splice site (A5SS), alternative 3’ splice site (A3SS) and a merge of the remaining event classes (other). **(D)** Heatmap shows changes in inclusion levels (ΔPSI) of cassette exons (rows) in the knockdown and overexpression conditions (columns). Events shown were quantified in all 10 comparisons and regulated in at least one of the two strongest knockdown and/or overexpression conditions. **(E)** Scatter plot shows *MST1R* exon 11 inclusion levels (ΔPSI) against HNRNPH protein levels. To test for cooperativity, a four parametric logistic curve (also known as Hill equation) was fitted to the data points (solid line). Hill coefficient (n_H_) and pseudo *R*^2^ (pR^2^) are indicated on the plot. **(F)** Histogram shows the distribution of Hill coefficient estimates for all HNRNPH-dependent cassette exons. Value ranges considered as cooperatively regulated (*n*_*H*_ ≥ 2) are indicated in blue (enhanced) and red (repressed); non-cooperative events are shown in white. **(G)** Scatter plots show inclusion levels (ΔPSI; quantified from RNA-seq) against HNRNPH protein levels for exons showing cooperatively enhanced (exon 5 of *PIP4K2C*; top left), repressed (exon 6 of *MYL6*; top right) and non-cooperative (exon 3 of *SNHG21* and exon 2 of *ZBTB24*; bottom) regulation. Data representation as in (E). **(H)** Genome browser view of cooperatively regulated cassette exon 5 of *PIP4K2C* (left) and exon 6 of *MYL6* (right). Shown are coverage and junction usage (PSI) of the strongest knockdown (46%), the knockdown control (100%), the overexpression control (100%), and the strongest overexpression (176%). Coverage is based on merged replicates and junction usage based on PSI estimates reported by MAJIQ, whereby usage of the skipping junction is the average of the PSIs reported for the two involved LSVs.

To identify HNRNPH-dependent splicing, we quantified alternative splicing changes using MAJIQ (Vaquero-Garcia et al. 2016) (**Figure 1A**). In total, we detected 1,828 alternative splicing events that significantly changed across the HNRNPH titration compared to Ctrl (measured as changes in relative junction usage, delta percent selected index [ΔPSI] > 0.025, probability > 0.9; **Figure 1C, Figure S1D, Supplementary Table S2**). The majority of regulated events were cassette exons (CE, 615), followed by multi-exon skipping (MES, 422) and intron retention (IR, 289) events, with splicing changes reaching up to 59% in either direction (**Figure 1C, Figure S1D**). In line with HNRNPH’s primarily repressive role, 71.8% of regulated CE showed reduced inclusion with rising HNRNPH levels (**Figure 1D**).

To examine cooperative regulation, we generated dose–response curves and fitted Hill equations for all regulated CE, IR, and alternative last exon (ALE) events (see Methods) (Braun et al. 2018). Most events achieved a high goodness-of-fit (331 CE [74.0%], 257 IR [93.1%], and 60 ALE [84.5%] with pseudo R^2^ ≥ 0.75 between best model fit and data; **Figure S2A, Supplementary Table S3**).

In line with previous semi-quantitative reverse transcription (RT)-PCR measurements (Braun et al. 2018), the dose–response curve for *MST1R* exon 11 from the RNA-seq data showed strong cooperative repression (Hill coefficient [n_H_] = 10.4; **Figure 1E**). Unexpectedly, almost all splicing changes were regulated cooperatively by HNRNPH: These included 302 cooperatively regulated CE (90.9%), mostly repressed by HNRNPH (n_H_ ≥ 2), with some showing particularly sharp responses, such as *MYL6* exon 6 (n_H_ = 25.3; **Figure 1F–H, S2B**). Similarly, 241 IR events (93.8%) and 56 ALE (93.3%) were cooperatively regulated by HNRNPH (**Figure S2B, C**).

To confirm splicing cooperativity by an independent method, we conducted RT-PCR experiments across the HNRNPH titration series for three CE. Consistent with the RNA-seq data, the dose–response curves from the RT-PCR measurements demonstrated strong cooperative regulation of *MYL6* exon 6 (n_H_ = 14.1) and *PIP4K2C* exon 5 (n_H_ = 11.4), while the control *SNHG21* exon 3 showed no cooperativity (n_H_ = 0.8; **Figure S2D**). Together, the results from the *in vivo* titration of cellular HNRNPH protein levels highlight a strong prevalence of cooperativity in HNRNPH-mediated splicing regulation.

### The positioning of HNRNPH binding determines the splicing outcome

To uncover the molecular mechanism behind HNRNPH-mediated regulation, we mapped HNRNPH binding sites in MCF7 cells using individual-nucleotide resolution UV crosslinking and immunoprecipitation (iCLIP2) (Buchbender et al. 2020; König et al. 2010). Across seven replicate experiments, we obtained a total of 178.4 million unique crosslink events for HNRNPH (**Supplementary Table S1**). Following peak calling by PureCLIP (Krakau, Richard, and Marsico 2017) and BindingSiteFinder (Mirko Brüggemann, Kathi Zarnack n.d.), we identified 468,452 high-confidence binding sites, predominantly located in the transcripts of protein-coding genes (85.7%; **Figure 2A**). Globally, 82.0% of all expressed protein-coding genes (transcripts per million [TPM] ≥ 1) harbored at least one binding site (**Figure S3A, B**). Within protein-coding transcripts, HNRNPH bound preferentially to intronic regions (88.6%; **Figure 2B**).

**Figure 2.**
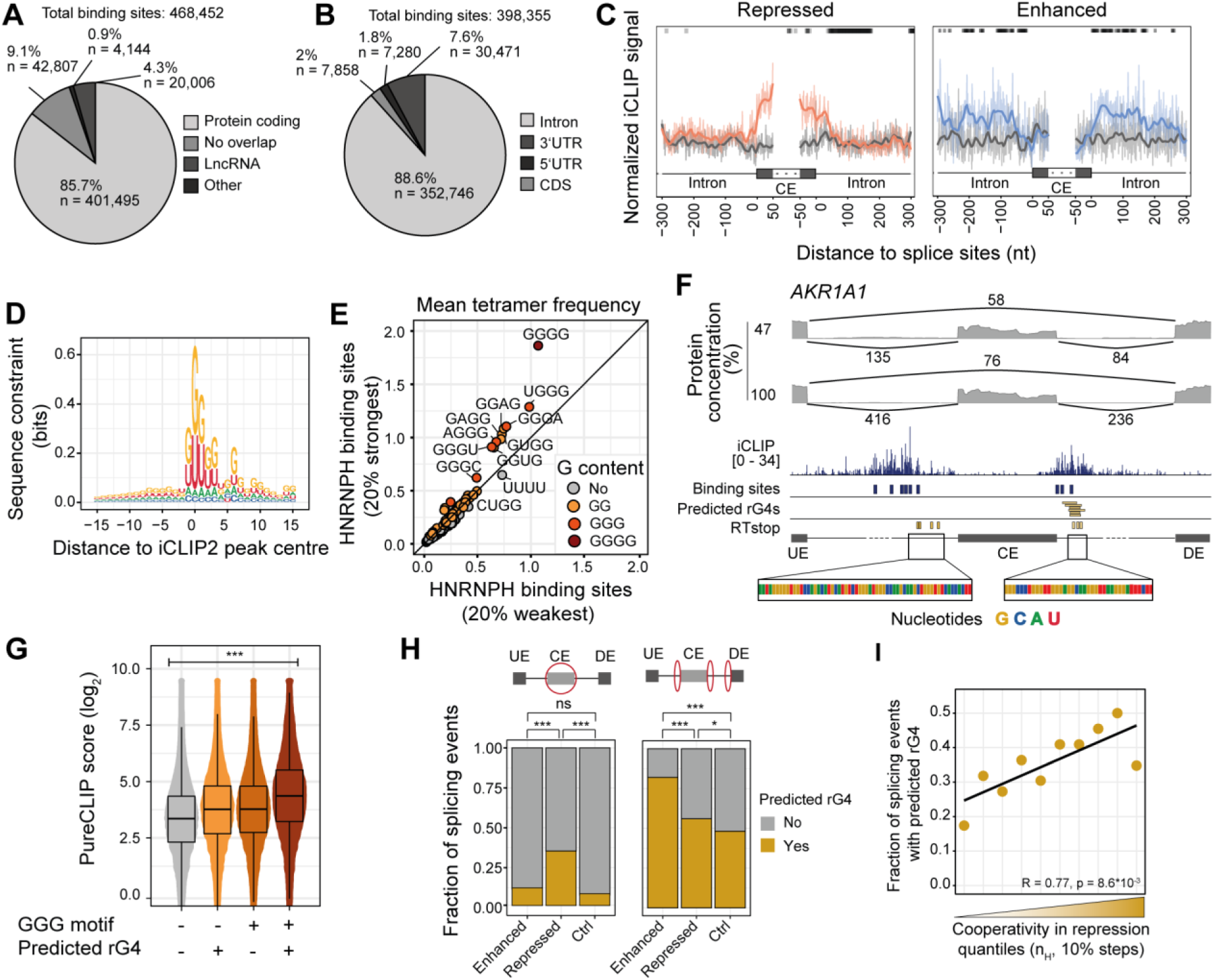
HNRNPH regulates target exons by direct binding in regions with high propensity for rG4 formation. **(A, B)** Pie charts show the percentage of HNRNPH binding sites overlapping with selected biotypes (A) and features of protein-coding genes (B). Numbers of binding sites considered for the pie charts are indicated. **(C)** Meta profiles of cooperatively repressed (left) and enhanced events (right) show normalized iCLIP2 signal at cassette exon (CE), and the flanking introns. The signal at cooperatively regulated events (red and blue lines) is compared against matched control (non-regulated) events (grey lines) and positions with a significant signal difference are indicated at the top of each panel. Raw signal and loess smoothing are shown. **(D)** Logo plot summarizes sequence composition of the 468,452 HNRNPH binding sites in a ±15-nt window aligned at the center of the binding sites. **(E)** Scatter plot shows the mean tetramer frequency in a ±25-nt window around the center of the 20% strongest and weakest binding sites. Tetramers are colored according to the number of consecutive Gs. **(F)** Genome browser view of CE and flanking introns in *AKR1A1* gene. Shown are coverage and read counts of the strongest knockdown (46%), the knockdown control (100%), as well as the raw iCLIP2 signal (shown range: 0–34 crosslink events) of HNRNPH binding, called binding sites, predicted rG4s and RTstop peaks. **(G)** Boxplots show the strength (log_2_-transformed PureCLIP score) of binding sites, which were divided into four groups based on the presence of a GGG motif in the binding site and whether the binding site overlaps with a predicted rG4. Statistical significance was assessed using a Kruskal–Wallis test across all groups, \*\*\**p* < 0.001 **(H)** Stacked bar charts show the distribution of cooperatively enhanced, repressed, and control splicing events overlapping with predicted rG4s (yellow) or not overlapping (gray), for the alternative exon (left panel) or its flanking introns (right panel). Red ellipses highlight the regions considered for overlaps with predicted rG4s. **(I)** Distribution of splicing events with predicted rG4 across ten quantiles based on their Hill coefficient (n_H_). Pearson correlation coefficients (*R*) and the associated *p*-value is indicated.

When integrating the iCLIP data with the observed splicing changes, we found that, consistent with previous studies (Chen, Kobayashi, and Helfman 1999; Markovtsov et al. 2000), HNRNPH preferentially bound within the cassette exon for HNRNPH-repressed CE, whereas it bound in the flanking introns of enhanced CE (**Figure 2C**). Notably, HNRNPH binding sites were more abundant at cooperatively regulated exons compared to control exons (**Figure S3C**), suggesting that cooperative regulation may arise from multivalent HNRNPH binding at these exons.

### rG4s are enriched at HNRNPH binding sites and cooperatively regulated exons

To elucidate the RNA binding specificity of HNRNPH, we investigated the sequence context of its binding sites. In line with its known sequence preference (Caputi and Zahler 2001), we observed an accumulation of G at the center of the HNRNPH binding sites, with G-rich tetramers increasing with binding site strength (**Figure 2D, E**). The G enrichment extended into the surrounding sequence, with recurrent G pileups within 15 nucleotides (nt) downstream of the binding sites, suggesting a potential RNA G4-quadruplex (rG4) formation (**Figure 2D**). To explore this, we used pqsfinder (Hon et al. 2017) which predicted putative rG4s at 121,321 HNRNPH binding sites (± 50 nt around the binding site center). This concurrence is exemplified at the cooperatively enhanced *AKR1A1* exon 7, where several predicted rG4s overlap with the HNRNPH binding sites (**Figure 2F**). Importantly, HNRNPH showed stronger binding in the presence of a predicted rG4 than at an isolated GGG motif (**Figure 2G**). Together, these observations suggest that rG4 structures enhance HNRNPH binding across thousands of binding sites in the transcriptome.

We compared the occurrence of predicted rG4s within regulated CE with control CE that were not regulated by HNRNPH. Interestingly, the number of predicted rG4s was significantly higher at cooperatively repressed exons, with 35.4% (79 out of 223) containing at least one predicted rG4, compared to 11.5% of cooperatively enhanced (9 out of 78) and 7.7% of control exons (89 out of 1,151; **Figure 2H**, left, **Figure S3D**). Furthermore, the presence of predicted rG4s correlated with the degree of cooperativity in repression, such that exons with steeper dose–response curves harbored more predicted rG4s (**Figure 2I**). Overall, the *in silico* predictions suggest that rG4 structures influence HNRNPH binding and contribute to cooperativity in splicing regulation.

### HNRNPH binding sites fold into RNA G-quadruplexes

To determine whether the *in silico* predicted rG4s fold in real transcripts, we used reverse transcriptase stop (RTstop) profiling (Guo and Bartel 2016). The method relies on the observation that rG4s and other strong RNA secondary structures impede reverse transcription (RT), resulting in truncated cDNAs that terminate at the RNA folds (**Figure S4A**). We adapted the RTstop protocol to a pool of 11,576 RNA *in vitro* transcripts (IVTs), representing 2,894 different transcript regions transcribed from a designed oligonucleotide library. Each 200-nt DNA oligonucleotide contained 146 nt of transcript sequence flanked by a T7 promoter for *in vitro* transcription and an L3 linker sequence for subsequent library preparation (**Figure S4A**). To comprehensively cover a wide range of HNRNPH binding regions, 1,340 HNRNPH binding sites with a predicted rG4 (BS+rG4) and 1,409 without rG4 (BS-only) were included in the oligonucleotide library (**Supplementary Table S4**). Additionally, we included 145 regions with predicted rG4s but no apparent HNRNPH binding (rG4-only). Each region was represented by four oligonucleotides with distinct barcodes to ensure robust and reproducible rG4 detection.

To distinguish rG4s from other RNA structures, the RT reaction was carried out in buffers containing either KCl or NaCl, with KCl stabilizing rG4s (Varshney et al. 2020) (**Figure S4A, B**). As an additional control, IVTs were also transcribed in the presence of the GTP analog 7-deaza-GTP (7dGTP), which cannot form rG4s via the Hoogsten interface but does not interfere with HNRNPH binding to linear RNA according to the published crystal structure (Dominguez et al. 2010). The experiment was performed in three replicates for each condition, with an average of 11.6 million reads per sample (**Supplementary Table S1**). The resulting RTstop signal showed characteristic pileups of read starts that were sensitive to the ion conditions. This is exemplified by the IVT harboring the HNRNPH binding site at *AKR1A1* exon 7 that showed three pileups present with KCl but absent with NaCl, suggesting that overlapping rG4s can form in this RNA (**Figure 3A**, rows 3 and 4). That these are indeed rG4s was further supported by their absence in the 7dGTP samples, regardless of the ion condition (**Figure 3A**, rows 5 and 6).

**Figure 3.**
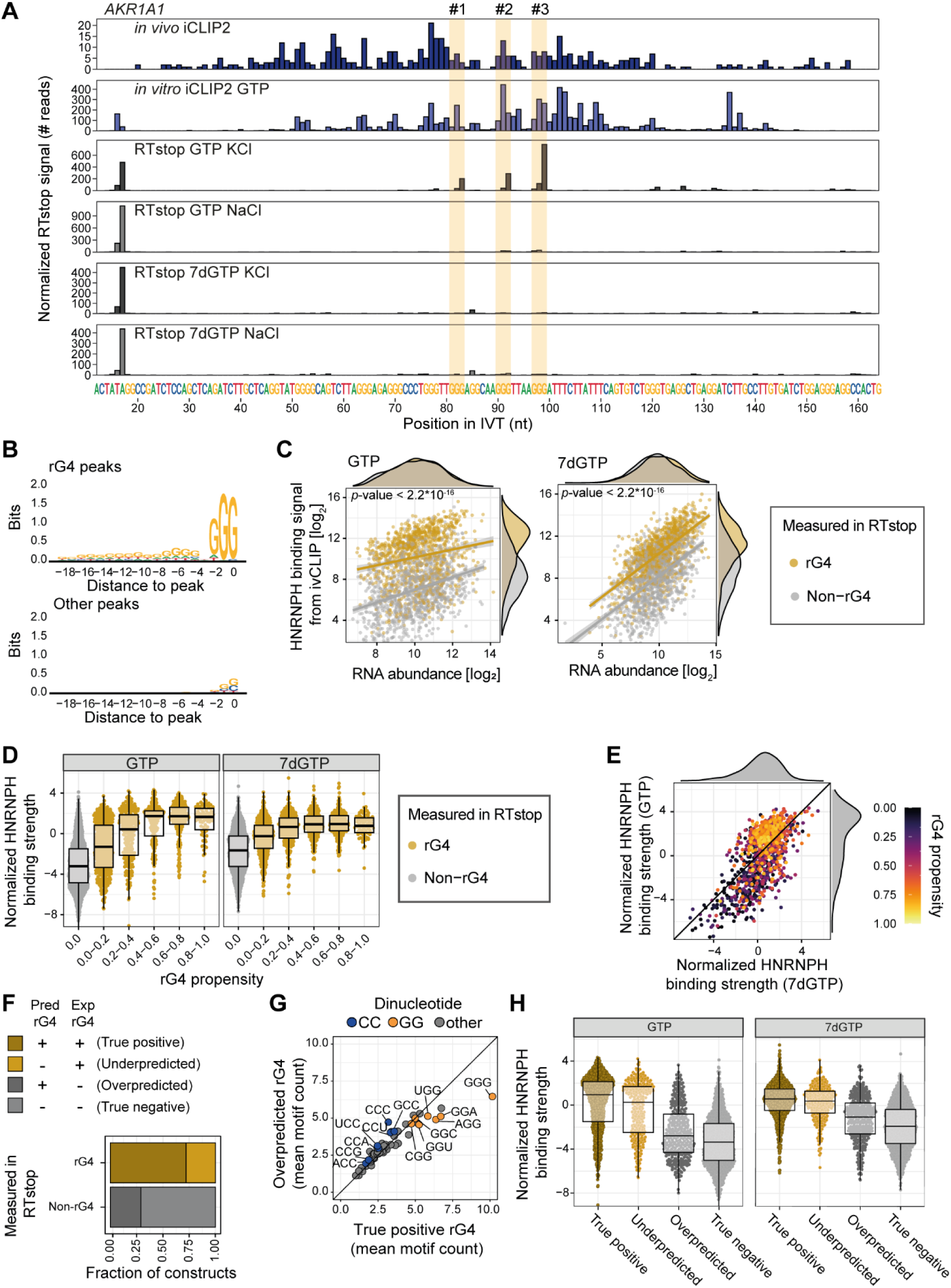
HNRNPH binding sites fold into RNA rG4 *in vitro*. **(A)** Genome browser view of an exemplary IVT (*AKR1A1*, chr1:45,568,657–45,568,802) showing changes in RTstop reads at selected GGG motifs (highlighted in yellow boxes) in conditions that stabilize (KCl) and destabilize (NaCl, 7dGTP) rG4. Reads from *in vivo* and *in vitro* HNRNPH iCLIP for selected region are also shown (rows 1 and 2). **(B)** Logo plot showing sequence in the 20-nt window from rG4 peak, i.e., responsive to the change in ion conditions, (upper plot) and other peaks (bottom). **(C)** Scatter plots show the correlation between RNA abundance (log₂) and HNRNPH1 binding signal (log₂) measured by *in vitro* iCLIP2 in the presence of either GTP (left) or 7dGTP (right). Each point represents a construct measured in the RTstop assay and is color-coded based on rG4 status (rG4, yellow; non-rG4, gray). Marginal density plots indicate the distribution of RNA abundance and HNRNPH binding levels. *P* value from two-sided Wilcoxon rank sum test. **(D)** Boxplots show the normalized HNRNPH binding strength across different rG4 propensities, comparing conditions with GTP (left) and 7dGTP (right). Data points for rG4 constructs (yellow) and non-rG4 constructs (gray) are displayed with violin plots indicating the distribution of binding strength. **(E)** Scatter plot with marginal density plots shows correlation between normalized HNRNPH binding strength with GTP (y-axis) and 7dGTP (x-axis). rG4 propensity is indicated by color scale. **(F)** Bar plot shows the fraction of constructs predicted to contain rG4s (darker shade) for both rG4 and non-rG4 regions measured in RTstop. **(G)** Correlation of trimer frequency between predicted rG4s but measured non-rG4s (y-axis; overpredicted) and predicted and measured rG4s (x-axis; true positive). Trimers are color-coded according to the consecutive dinucleotides. **(H)** Boxplots show the normalized HNRNPH binding strength for predicted and non-predicted rG4s, categorized based on RTstop measurements as either rG4 or non rG4, comparing conditions with GTP (left) and 7dGTP (right). Data represented as in (F).

A quantitative comparison between the KCl and NaCl conditions identified 8,986 rG4s in 1,060 tested transcript regions (false discovery rate [FDR] < 0.05), whereas 1,109 regions were devoid of rG4s. The remaining IVTs showed an intermediate result and were excluded from further analysis (see Methods). Importantly, we detected at least one rG4 in 687 out of 982 (70.0%) predicted BS+rG4 regions that could be assigned. In contrast, only 25.1% (250 out of 996) of predicted BS-only constructs exhibited measurable rG4 formation (**Supplementary Table S5**). The intensity of the RTstop signal for the rG4s corresponded to the prominence of G-runs in the RNA sequence, indicating that this feature provides quantitative information on the degree of rG4 formation (**Figure 3B, Figure S4C**). These findings confirm that many HNRNPH binding sites show a strong propensity to fold into rG4s *in vitro*.

To directly assess how rG4 formation impacts HNRNPH binding, we performed *in vitro* iCLIP2 on the IVTs using recombinant HNRNPH1 protein. The IVT library was transcribed in the presence of either GTP or 7dGTP to compare HNRNPH1 binding to the same RNAs with or without rG4s, respectively (four replicates per condition; on average ∼20 million reads per sample; **Supplementary Table S1**). Intriguingly, rG4-containing RNAs received significantly stronger HNRNPH1 binding than non-rG4 RNAs (**Figure 3C**, left). Moreover, among the rG4-containing RNAs, HNRNPH1 binding increased progressively with the degree of *in vitro* rG4 formation (**Figure 3D, Figure S4D, E**). Notably, the difference between rG4-containing and non-rG4 was diminished when rG4 formation was prevented by 7dGTP (**Figure 3C, D**, **Figure S4D, E**), and rG4-containing RNAs showed significantly stronger HNRNPH1 binding in the GTP condition compared to the 7dGTP condition (**Figure 3E**). Overall, the presence of an rG4 enhances HNRNPH binding.

The high-throughput measurements allowed us to evaluate the accuracy of the *in silico* rG4 predictions across hundreds of sequences. Specifically, pqsfinder failed to detect rG4s in 294 (27.9%) of RNAs with an experimentally confirmed rG4s (false-negative; underpredicted), while incorrectly predicting rG4s in 322 (29.0%) of the rG4-devoid RNAs (false-positive, overpredicted; **Figure 3F**). Sequence analysis revealed an enrichment of cytosine (C)-rich triplets around the false-positive rG4 predictions (**Figure 3G**), suggesting that an excess of C may interfere with rG4 formation. A higher reliability of the experimentally determined rG4s over the *in silico* predictions was supported by the *in vitro* iCLIP2 results: RNAs with false-negative prediction clearly showed strong HNRNPH binding, similar to true-positive RNAs (**Figure 3H**). In contrast, RNAs false-positive prediction exhibited only weak HNRNPH binding, consistent with the lack of rG4 formation *in vitro*. These findings suggest that the *in silico* predictions capture the true rG4 formation only partially and support the accuracy of the experimental rG4 measurements. Altogether, we conclude that the RNA sequences at HNRNPH binding sites show a strong propensity to fold into rG4 structures which, in turn, enhances HNRNPH binding.

### rG4 unfolding establishes indirect cooperativity in HNRNPH binding

RBP binding to dynamic secondary structures opens the possibility for indirect cooperativity when binding alters the RNA’s structure (Becker et al. 2019). To investigate whether HNRNPH influences rG4 folding, we performed fluorescence resonance energy transfer (FRET) experiments. We designed a 23-nt rG4 RNA probe, derived from a published HNRNPH-bound rG4 in the *ARPC2* transcript (von Hacht et al. 2014), with a 5’ donor FAM fluorophore and a 3’ acceptor TAMRA fluorophore (**Figure 4A**). These fluorophores are brought into close proximity when the rG4 is formed, allowing the FAM donor to transfer its emission to the TAMRA acceptor, quenching the FAM signal (De Cian et al. 2007; Malina, Scott, and Brabec 2020). Indeed, FAM emission was significantly lower under rG4-stabilizing (KCl) compared to non-stabilizing (LiCl) conditions, confirming that the probe accurately quantifies rG4 formation *in vitro* (**Figure 4B, Figure S5A**). Importantly, when increasing concentrations of recombinant HNRNPH1 protein were added to the folded rG4s (KCl), FAM emission significantly increased up to 140% (**Figure 4B**), indicating rG4 unfolding. This supports previous reports showing that HNRNPH1 preferentially binds to unfolded G-runs (Vo et al. 2022) and suggests that HNRNPH binding prevents refolding by stabilizing the unfolded conformation.

**Figure 4.**
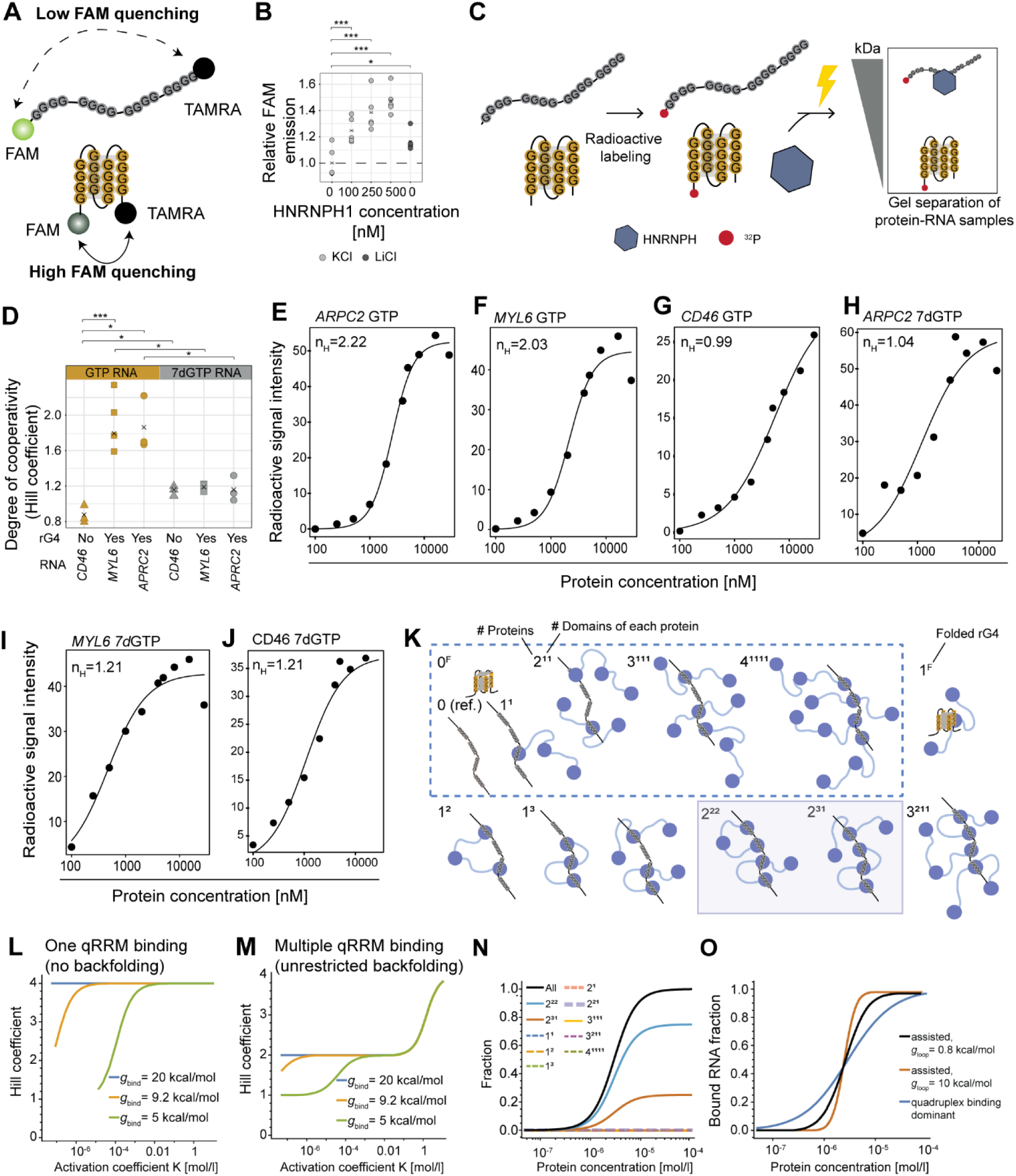
Recombinant HNRNPH1 binds cooperatively to rG4 RNAs *in vitro*. **(A)** Schematic illustration of the design of a 23-nt rG4 RNA probe with a 5’ donor FAM fluorophore and a 3’ acceptor TAMRA fluorophore that are brought in close proximity when the rG4 is formed. The excited donor FAM then transfers its emission to the acceptor TAMRA, thereby quenching the FAM emission signal (De Cian et al. 2007; Malina, Scott, and Brabec 2020). **(B)** Plot shows FAM emission of an rG4-forming FAM-TAMRA-labeled RNA oligonucleotide (shown in **Figure S5B**) in an rG4-disfavoring (LiCl) condition and with increasing concentrations of recombinant HNRNPH1 protein relative to the FAM emission of the RNA oligonucleotide in an rG4-favoring condition (KCl). FAM excitation was performed at 485 nm and FAM emission was measured at 520 nm. All experiments were performed in five individual replicates and to test for significant differences a two-sample t-test was performed. **(C)** Experimental design of *in vitro* titration experiments with recombinant HNRNPH1. RNAs were radioactively labeled and incubated with recombinant HNRNPH1, followed by UV crosslinking to stabilize the protein-RNA complexes. The formation of these complexes was quantified by separating them from unbound RNA based on size. **(D)** Plot shows the degree of cooperativity (Hill coefficient) for recombinant HNRNPH1 protein titration to three different RNA oligonucleotides with and without rG4 formation, *in vitro* transcribed with GTP (left) or 7dGTP (right). To test for significant differences of the Hill coefficients between GTP and 7dGTP conditions, a Welch two-sample *t*-test was performed. (**E–J**) Scatter plots show the signal intensity of the radioactively labeled RNA from the protein–RNA complexes against recombinant HNRNPH1 protein concentration in presence of GFP (E–G) and 7dGTP (H–J) for two examples. Hill equations were fitted to the scatter plots to test for cooperativity using the Hill coefficient (n_H_). **(K)** Possible binding modes with corresponding notation. The dashed box shows modes contributing to the simplified double backfolding model. **(L, M)** Hill coefficient vs. K (tuned by varying the folding energy) for extreme cases of no backfolding penalty (L) and unrestricted backfolding penalty (M). **(N)** Contribution of different binding modes to the total fraction of bound RNA for folding energy *g*_rG4_ = 23 kcal/mol, *g*_bind_ = 9.2 kcal/mol, and *g*_backfold_ = 0.8 kcal/mol. **(O)** Bound RNA fraction vs. protein concentrations for folding energy *g*_*rG*4_ = 23 kcal/mol and *g*_*bind*_ tuned such that *K* = 3 ⋅ 10^−6^mol/l for two cases of low and high backfold penalty *g*_*backfold*_ (giving *n*_*H*_ = 2 and *n*_*H*_ = 4, respectively), without and with assistance by a helper state 1^*F*^ with binding energy *g*_*rG*4,*bind*_ = 4 kcal/mol (red and blue lines). If *g*_*rG*4,*bind*_ is too strong, the cooperativity gets lost (black line corresponding to *g*_*rG*4,*bind*_ = 6.1 kcal/mol and 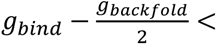 8 kcal/mol).

Next, we investigated whether rG4 unfolding facilitates cooperative HNRNPH binding to RNA. For this, we quantified RNA–protein complex formation in *in vitro* titration experiments with recombinant HNRNPH1 protein and *in vitro* transcribed RNAs (**Figure 4C**). In addition to a fragment from the *ARPC2* transcript (190 nt), we used an rG4-forming HNRNPH binding site from the HNRNPH-regulated *MYL6* exon 6 (71 nt), and a control HNRNPH binding site from *CD46* mRNA lacking an rG4 (112 nt; **Supplementary Table S6**). Circular dichroism spectroscopy confirmed that both *ARPC2* and *MYL6* RNAs formed parallel-stranded rG4s in the presence of KCl, whereas the *CD46* RNA displayed no rG4 (**Figure S5B**) (Randazzo, Spada, and da Silva 2013; Vorlíčková et al. 2012). Importantly, when the RNAs were incubated with increasing HNRNPH1 protein concentrations (100 nM–28.5 µM), we observed positive cooperativity in HNRNPH1 binding to the rG4-containing *ARPC2* and *MYL6* RNAs (mean n_H_ = 2.22 and 2.03; **Figure 4D–F, Figure S5C**). The Hill coefficient of n_H_ ∼ 2 indicates that two HNRNPH1 molecules conjointly bound, such that binding of the first protein made it easier for the second protein to bind. In contrast, the non-rG4 RNA *CD46* exhibited only non-cooperative binding in the absence of an rG4 (mean n_H_ = 0.99; **Figure 4D, G, Figure S5D**). To support that cooperativity depends on rG4 formation, we repeated the experiments using the same RNAs transcribed with 7dGTP to prevent rG4 formation. In the absence of rG4s, HNRNPH bound non-cooperatively to all three RNAs (**Figure 4D, H–J, Figure S5D**). Together, these results reveal that two HNRNPH molecules bind cooperatively to RNAs that form rG4 structures.

### Multiple qRRMs of HNRPNH participate in binding the unfolded rG4

Unfolding of the rG4 structure exposes four single-stranded G-runs, creating potential binding sites for up to four qRRMs. To explore possible binding states (**Figure 4K**), we developed a quantitative biophysical model that describes the system using three characteristic free energies: (i) the free energy difference between folded and unfolded rG4, denoted *g*_rG4_, (ii) the binding free energy for a single HNRNPH binding event, denoted *g*_bind_, and (iii) an entropic penalty for multi-qRRM binding which arises if an HNRNPH protein folds back to allow an additional qRRM of the same protein to bind, denoted *g*_backfold_. For simplicity, we treated the three qRRMs of HNRNPH as equivalent. Using the unfolded state as a reference, we derived the fraction of rG4s bound to at least one protein at equilibrium, the transition concentration (activity coefficient *K*) and the Hill coefficient. To analyze the behavior of the model, we first considered two extreme cases: In the first case, we allowed each HNRNPH protein to bind with only one qRRM (no backfolding, dashed box in **Figure 4K**)This resulted in a Hill coefficient of *n*_*H*_ = 4 for sufficiently large binding and folding energies (**Figure 4L**), thus diverging from the experimentally observed Hill coefficient of *n_H_* ∼ 2. In the second case, we introduced unrestricted backfolding (i.e., a backfolding penalty *g*_backfold_ = 0), resulting in *n*_*H*_ = 2 for a broad parameter range (**Figure 4M**). Notably, an *n*_*H*_ ≤ 2 was achieved for all activation coefficients *K* < 0.1 mol/l, independent of the binding energy, *g*_bind_, indicating that backfolding and binding of more than one qRRM generally occur. Further inspection revealed that systems with *n*_*H*_ ≈ 2 were dominated by two binding states, namely 2^31^ and 2^22^ (2^31^ referring to two HNRNPH molecules, binding with 3 and 1 qRRM, respectively; **Figure 4K**), each involving two backfolding events such that all four RNA binding sites were occupied (see below).

Next, we used the experimental data to refine the model. First, we set an activity coefficient *K* ≈ 3 · 10^−6^ mol/l, corresponding to the protein concentration at the transition (**Figure 4E–J**) This constrained the free energy parameters. For *K* = 3 · 10^−6^mol/l, a reduced model incorporating only the two dominant binding states (2^31^ and 2^22^, termed “double-backfolding model”) described the data similarly to the full model for all *g*_backfold_ < 7 kcal/mol (**Figure S5E**). For higher *g*_backfold_, the Hill coefficient increased to *n*_*H*_ = 4, indicating the suppression of the two dominant binding states due to a stronger backfolding penalty (**Figure S5F**). To determine *g*_backfold_, we estimated the rG4 folding energy as *g*_rG4_ ≈ 23 kcal/mol (measured by (Yu et al. 2009), and the HNRNPH binding energy as *g*_bind_ = 9.2 kcal/mol, derived from the molecular dissociation constant *K*_*d*_ = 0.2 · 10^−6^ mol/l measured for binding of the *Drosophila* HNRNPF/H ortholog Glorund to G-tract RNAs (Tamayo et al. 2017). Inserting these values yielded a backfolding penalty of *g*_backfold_ = 0.8 kcal/mol. The calculation indicated that the binding states were mostly dominated by the state 2^22^, followed by 2^31^ as the second-most probable state (**Figure 4N**). Thus, we propose that binding involves two HNRNPH proteins, which in total contact the RNA through two qRRM domains, thereby occupying all four binding sites.

Our FRET experiments suggested that initial HNRNPH binding to the folded rG4 results in unfolding (**Figure 4B**). To test this in our model, we considered the impact of a helper state, 1^*F*^, where HNRNPH binds to the folded rG4 with binding energy *g*_rG4,bind_, to facilitate unfolding. We found that if *g*_rG4,bind_ was less than 4 kcal/mol, the Hill coefficient did not change. However, if the binding energies to the folded and unfolded rG4 became comparable, binding to the folded rG4 began to dominate, and cooperative binding was lost (**Figure 4O**). This suggests that at sufficiently low binding energy *g*_rG4,bind_, HNRNPH binding to the folded rG4 the dynamics of the system and facilitates rG4 unfolding, thereby helping to overcome the potential barrier, without affecting the cooperativity. Altogether, our biophysical model indicates that cooperative binding of HNRNPH to rG4s involves backfolding of the protein to bind with multiple qRRMs to the exposed G-runs, thereby stabilizing the unfolded RNA structure.

### rG4s at HNRNPH binding sites are required for cooperative splicing regulation

Our *in vitro* experiments and theoretical models revealed that an RNA’s ability to form an rG4 indirectly facilitates cooperative HNRNPH binding. To test whether this mechanism supports the switch-like splicing regulation by HNRNPH *in vivo*, we conducted a series of minigene experiments, combining *HNRNPH* KD with binding site mutations. The minigene reporter included *AKR1A1* exon 7 together with its flanking introns and exons (**Figure 5A, Supplementary Table S7**). This CE is enhanced by HNRNPH (Dardenne et al. 2014; Zhang, Harvey, and Cheng 2019), displays strong cooperativity in its dose-response (**Figure 5B**) and contains HNRNPH binding sites with overlapping rG4s in both flanking introns (**Figure 2F**). By gradually reducing HNRNPH levels via KD in MCF7 cells, we confirmed that CE inclusion of the *AKR1A1* wildtype (wt) minigene was cooperatively enhanced by HNRNPH, mirroring the endogenous *AKR1A1* exon 7 (**Figure 5C, Figure S6A**).

**Figure 5.**
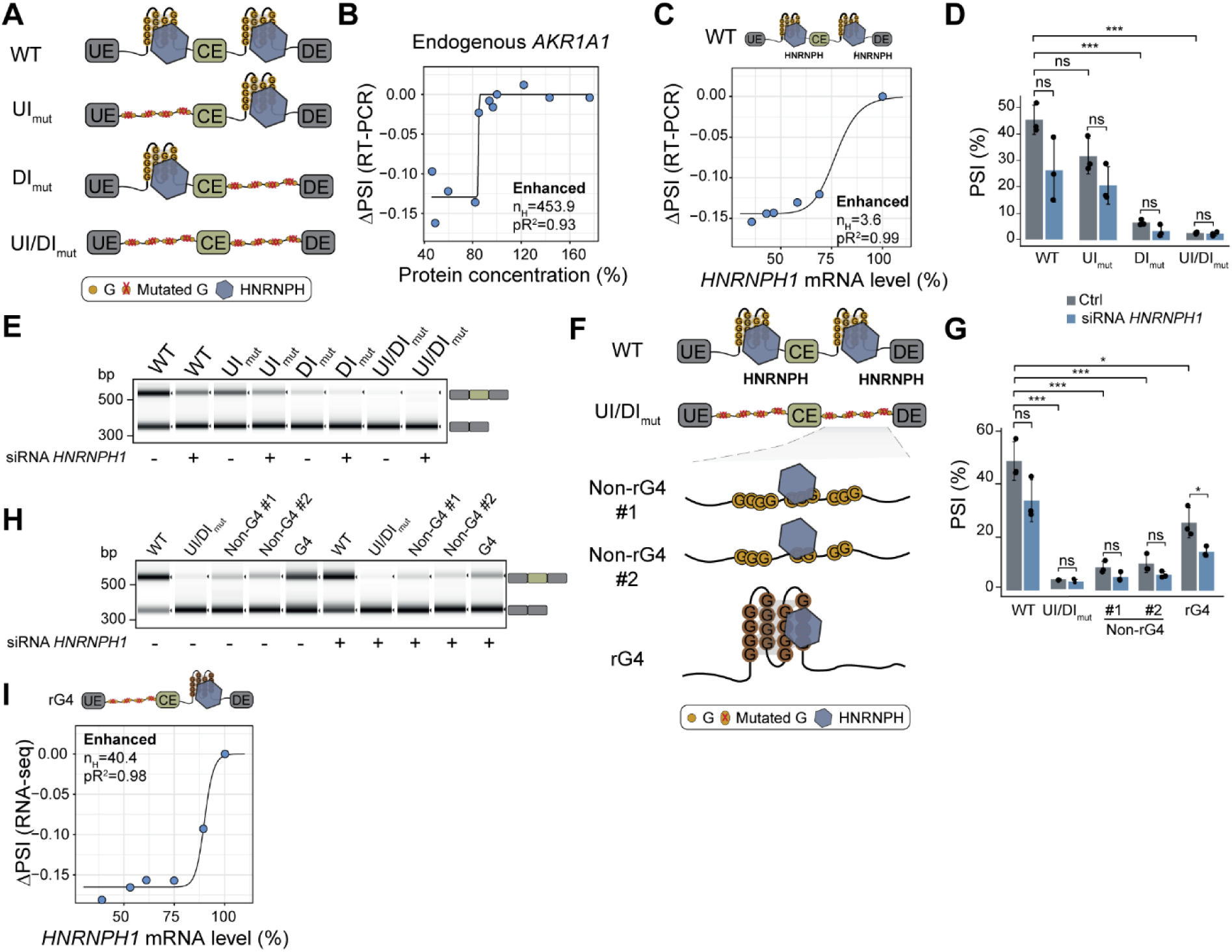
AKR1A1 minigene splicing depends on an rG4-forming HNRNPH binding site. **(A)** Schematic illustration of mutational constructs of the AKR1A1 minigene. The wildtype (WT) AKR1A1 minigene contains two rG4-forming HNRNPH binding sites, one in the upstream intron (UI) and one in the downstream intron (DI) of exon 7. Both HNRNPH binding sites were mutated separately (UI and DI constructs) and one AKR1A1 minigene construct carrying both mutations (UI/DI construct) was generated. **(B)** Scatter plot shows the splicing change (ΔPSI) of *AKR1A1* exon 7 against HNRNPH1 protein level in MCF7 cells. A Hill equation was fitted to the scatter plot to extract the Hill coefficient (n_H_) that was used to test for cooperativity. **(C)** Scatter plot shows the splicing change (ΔPSI) of the WT AKR1A1 minigene against HNRNPH1 protein level in MCF7 cells. Data represented as in B. **(D)** Bar plot shows the percent spliced-in (PSI) of *AKR1A1* exon 7 for the WT and mutational AKR1A1 minigene constructs described in (A), in control and *HNRNPH1* depleted condition. Mean and the error bars indicating the standard deviation of three biological replicates are shown. **(E)** Representative capillary electrophoresis image of semiquantitative RT-PCR results for *AKR1A1* splicing analyses shown in (D). **(F)** Schematic illustration of the construct design for new HNRNPH binding sites in the downstream intron of the AKR1A1 minigene construct UI/DI. Two constructs with HNRNPH binding sites without rG4s (non-G4 #1 and #2) and one new construct with an HNRNPH binding overlapping with an rG4 (rG4) were generated. **(G)** Barplot shows the percent spliced-in (PSI) of *AKR1A1* exon 7 for the WT and mutational AKR1A1 minigene constructs described in (F). Data represented as in (D). **(H)** Representative capillary electrophoresis image of semiquantitative RT-PCR results for *AKR1A1* splicing analyses shown in (G). **(I)** Scatter plot shows the splicing change (ΔPSI) of the rG4 AKR1A1 minigene against HNRNPH1 protein level in MCF7 cells. Data represented as in (B).

Next, we mutated both HNRNPH binding sites by substituting the middle G in each GGG trimer or the two middle G in each GGGG tetramer in both the upstream and downstream intron (UI/DI), which abolished any HNRNPH-dependent splicing regulation (**Figure 5A, D, Figure S6B, Supplementary Table S7**). Disrupting only the HNRNPH binding site in the upstream intron (UI) did not impact regulation, whereas mutating the downstream site (DI) significantly reduced CE inclusion under control conditions, along with a decrease in splicing regulation upon *HNRNPH* KD (**Figure 5A, D, E, Figure S6C, D**). Consistent with previous findings (Dardenne et al. 2014), our results thus indicate that the downstream HNRNPH binding site is important for *AKR1A1* exon 7 inclusion.

To assess whether the overlap with an rG4 is necessary for HNRNPH’s regulation of *AKR1A1* splicing, we replaced the downstream binding site with two different non-rG4-forming HNRNPH binding sites (as confirmed by our RTstop profiling; Non-rG4 #1 & 2) or with another rG4-forming HNRNPH binding site (**Figure 5F, Figure S6E, Supplementary Table S7**). Neither of the two non-rG4 binding sites restored HNRNPH-mediated splicing regulation in the *AKR1A1* minigene, suggesting that not just cooperativity but also overall HNRNPH regulation depends on rG4 formation (**Figure 5G, H, Figure S6E**). In contrast, inserting the rG4-forming HNRNPH binding site in the downstream intron reinstated the cooperative splicing regulation by HNRNPH (**Figure 5G–I, Figure S6E**).

In conclusion, our *AKR1A1* minigene reveals that robust, cooperative regulation of *AKR1A1* exon 7 splicing by HNRNPH depends on an intronic rG4-forming HNRNPH binding site positioned downstream of the alternative exon.

### Cooperativity is amplified in multi-step splicing reactions

Our biophysical model, along with the *in vitro* data, suggests Hill coefficients of *n*_*H*_ ≈ 2 for cooperative HNRNPH binding, albeit we observe substantially higher cooperativity in splicing regulation *in vivo*. To understand the link between cooperative HNRNPH binding and the more pronounced effect on alternative splicing, we employed kinetic modeling of splicing decision making. In our model, we conceptualized splicing regulation as a series of consecutive biochemical events (**Figure 6A**): in addition to cooperative HNRNPH binding to pre-mRNA (step 1), we considered that once bound, HNRNPH influences the sequential multi-step spliceosome assembly on nearby splice sites (step 2). Finally, spliceosome binding patterns on splice sites determine splicing decisions about exon inclusion or skipping (step 3).

**Figure 6.**
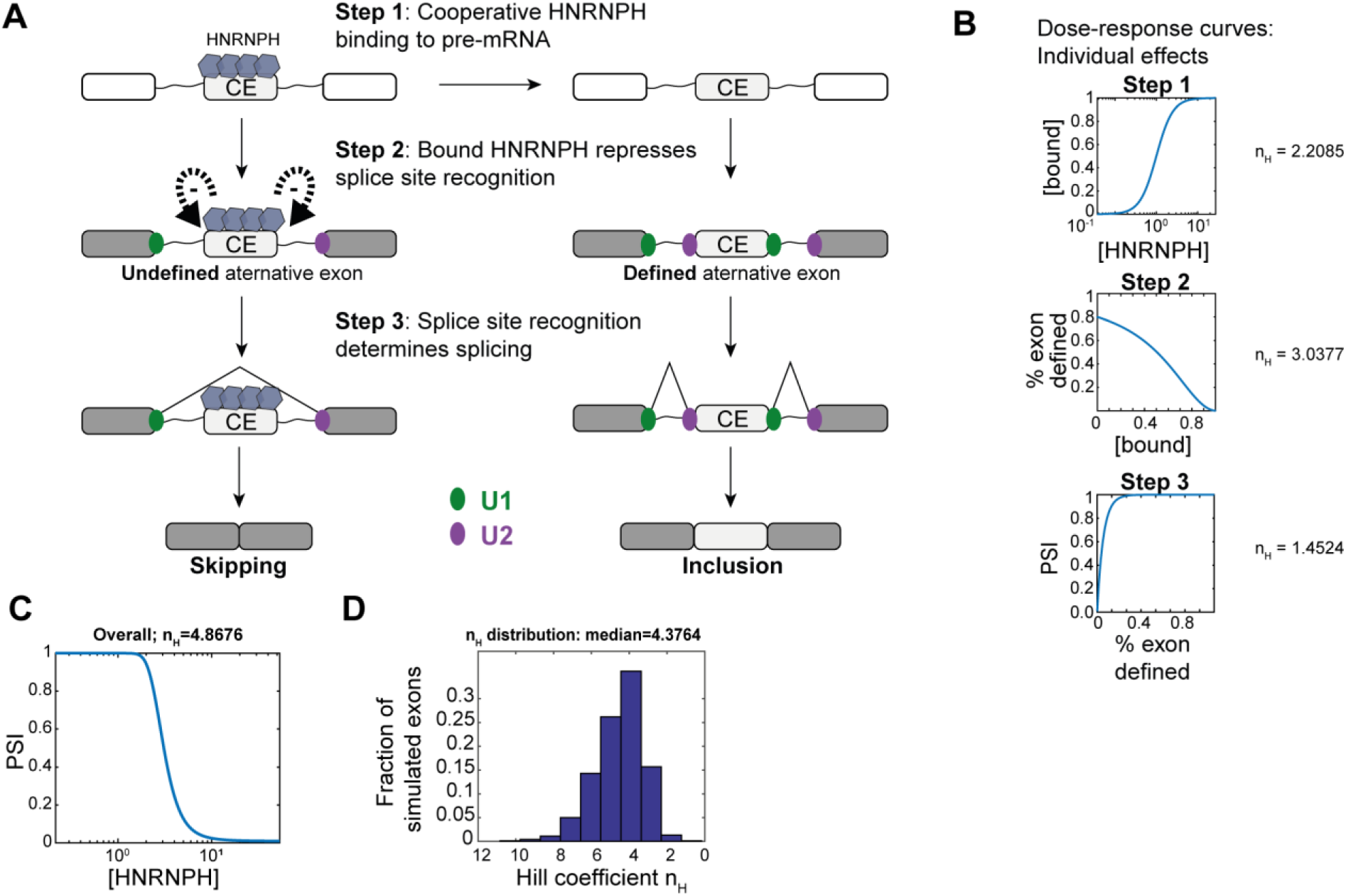
Amplification of switch-like behavior in a splicing-regulatory cascade. **(A)** Modeling HNRNPH-mediated splicing regulation as a cascade of consecutive biochemical steps. After cooperative HNRNPH binding to pre-mRNA in or nearby the alternative exon (step 1) the bound HNRNPH aids the recruitment of pioneering spliceosome subunits U1 and U2 to splice sites (step 2). Based on U1 and U2 binding patterns, splicing decisions are made (step 3), i.e., the alternative cassette exon is included or skipped depending on whether it is defined (U1&U2 bound) or not. **(B–D)** Moderate Hill coefficients at every level can result in highly switch-like behavior of the overall splicing-regulatory cascade. **(B)** Hill coefficients of *n*_*H*_ > 1 can be observed at every cascade level assuming the steady state equations and parameter given in the Supplemental Material due to cooperative HNRNPH binding (top), multistep spliceosome binding regulation by bound HNRNPH (middle, **Figure S7A**) and the co-transcriptional nature of splicing decision making (bottom, **Figure S7B**). **(C)** The overall cascade response, linking the PSI to the HNRNPH concentration, can be highly switch-like, with *n*_*H*_ = 4.8676 for the chosen parameter values (**Supplemental Material**). **(D)** A broad distribution of overall Hill coefficients n_H_ is observed when randomly sampling all kinetic parameters of the splicing cascade model to reflect the heterogeneity of individual exons. The histograms show Hill coefficients obtained for 1000 simulation runs (underlying dose–response curves shown in **Figure S7C**).

Each level in this splicing cascade exhibits sigmoidal dose–response behavior with moderate Hill coefficients. For step 1, we assumed that HNRNPH binding to pre-mRNA is cooperative as observed *in vitro* (**Figure 3**). In the subsequent spliceosome assembly, a Hill coefficient *n*_*H*_ > 1 can arise if we assumed that HNRNPH regulates the process at multiple levels, e.g. during both U1 and U2 binding (step 2). Finally, due to time delays in transcription-coupled splicing, the exon-inclusion decision (PSI) can depend in a sigmoidal fashion on the degree of spliceosome binding (step 3; **Figure S7A, B**, see **Supplemental Material**). To model the steady-state behavior of the splicing cascade, we used three algebraic equations, with the output from each stage feeding into the next. Previous studies have shown that signaling cascades with a similar architecture can amplify signals and yield highly switch-like responses (Ferrell and Ha 2014). Specifically, when each level is described by the sigmoidal Hill equation (with Hill coefficients *n*_*H*1_ − *n*_*H*3_), the cascade’s overall Hill coefficient can maximally become the product of the individual coefficients (*n*_*H*_ = *n*_*H*1_ × *n*_*H*2_ × *n*_*H*3_). Thus, even moderate cooperativity (*n*_*H*_ = 2) at each level can yield a very pronounced overall switch-like behavior (*n*_*H*_ ∼ 8).

In the splicing model, the resulting Hill coefficient depends on specific kinetic parameters, including HNRNPH affinity, spliceosome sensitivity to HNRNPH-mediated regulation, and inherent splice site strength. For a specific parameter combination, illustrated in **Figure 6C**, we observe an overall Hill coefficient of *n*_*H*_ = 4.9. To reflect the variability in splicing parameters across exons in the human transcriptome, we ran simulations with randomly sampling all parameter values in the model, assuming log-normal distributions (see **Supplemental Material**). Consistent with our experimental observations (**Figure 1F**), the simulated population of exons showed a wide distribution of Hill coefficients (**Figure 6D, Figure S7C**). Notably, larger Hill coefficients (*n*_*H*_ > 10), which were occasionally seen experimentally, were also achievable in our model. Overall, the simple splicing cascade model can explain why *in vivo* Hill coefficients of HNRNPH-mediated splicing regulation differ across exons and reach substantially higher than those observed for *in vitro* HNRNPH binding.

### HNRNPH-dependent splicing patterns discriminate breast cancer subtypes

Our experiments revealed that rG4s are critical for HNRNPH binding and splicing regulation in MCF7 cells. To test whether this is affected in disease, we screened genomic single nucleotide variants in the Penn Medicine BioBank (PMBB), including breast cancer patients from European (*n* = 29,362) and African (*n* = 10,217) ancestry. Using G4mer (Zhuang et al. 2024), we predicted 13,473 rG4-disrupting variants. Notably, 8 of rG4-disrupting variants were significantly enriched in the breast cancer samples, including two that neighbored HNRNPH-regulated alternative splicing events (**Figure 7A, Figure S8A**). This is exemplified by the variant rs139161442 (chr11:67310190:G:T) in the gene *SSH3* which lies in an exon with alternative 5’ splice sites responsive to HNRNPH regulation. Together, these observations indicate that variants changing individual HNRNPH–rG4 interactions could contribute to breast cancer phenotypes.

**Figure 7.**
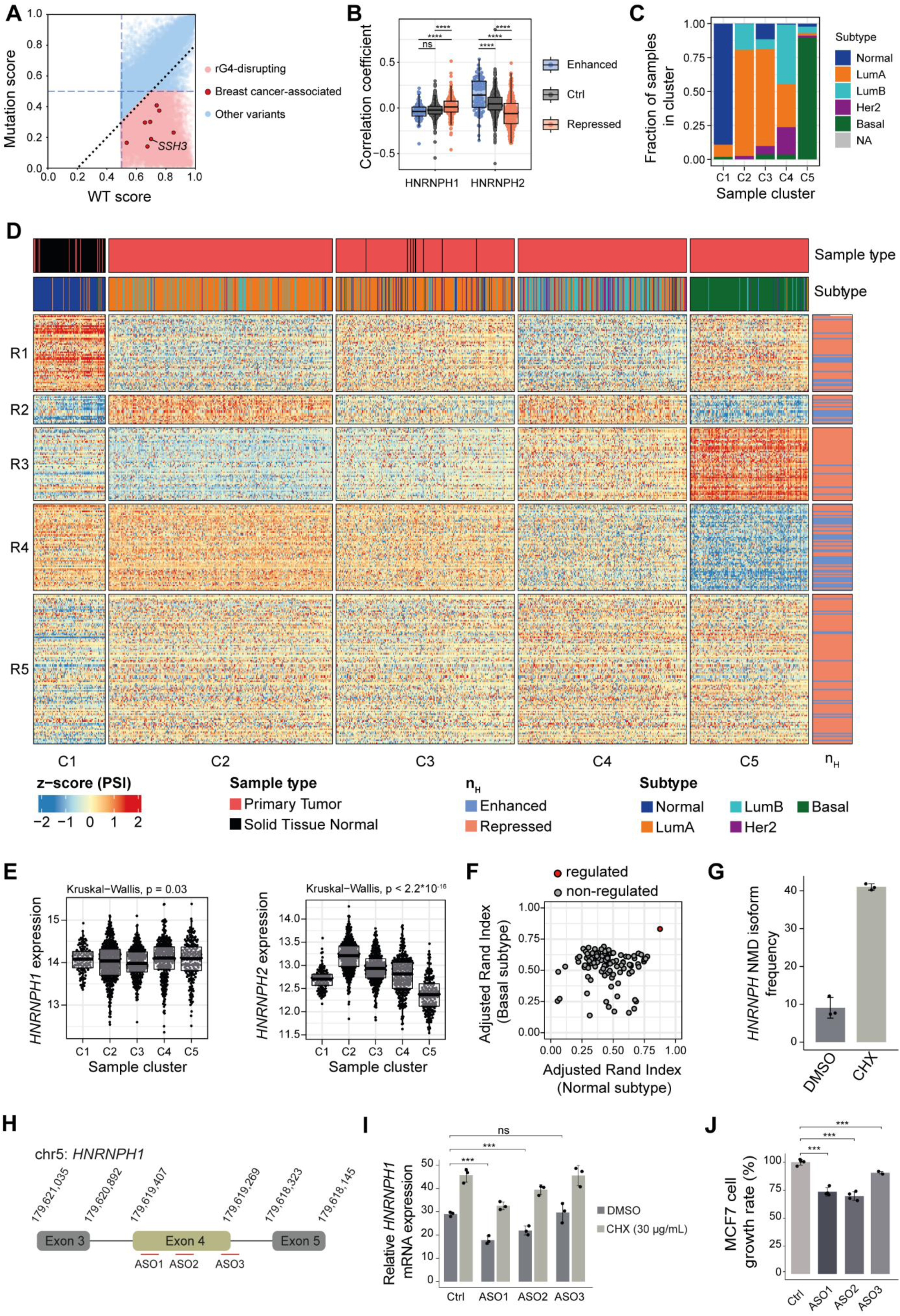
HNRNPH-dependent splicing discriminates breast cancer subtypes, and its downregulation inhibits cell growth. **(A)** Scatter plot shows correlation between *wildtype* (WT) and mutation score for each event. rG4-disrupting events are shown in pink, rG4 non-disrupting events in blue, and breast cancer–associated variants are highlighted in red. Events below the diagonal indicate greater rG4 disruption. **(B)** Correlation coefficient between HNRNPH-regulated CE (cooperatively enhanced in blue, cooperatively repressed in red, and control in gray) and expression of *HNRNPH1* and *HNRNPH2* across patient samples. **(C)** Stacked bar plot shows fractions of tumor subtypes in each column cluster. **(D)** Heatmap shows relative inclusion levels of HNRNPH-regulated CE (*n* = 302) across cancer (*n* = 1,069) and adjacent normal (*n* = 108) samples from the TCGA breast adenocarcinoma (BRCA) cohort. Five clusters of CE splicing (R1–R5) and patient samples (C1–C5) were obtained by *k*-means clustering of *z*-score-transformed PSI values. Annotations indicate sample type, molecular subtype, and splicing regulation in MCF7 cells (enhanced/repressed). **(E)** Plots show vst-transformed (variance-stabilizing transformation) *HNRNPH1* (left) and *HNRNPH2* (right) expression changes across five column clusters representing tumor subtypes from C1 (WT) to C5 (most severe basal subtype). *P* values from Kruskal-Wallis test. **(F)** Scatter plot shows correlation between adjusted rand index between normal and basal subtype. Regulated events are highlighted in red. **(G)** Bar plot shows *HNRNPH1* exon 4 skipping isoform frequency in control (DMSO) and NMD-inhibiting (cycloheximide, CHX) condition in MCF7 cells. Data points representing biological replicates and error bars representing standard deviation are shown. **(H)** Schematic illustration of the locations of ASOs designed to shift splicing towards the NMD-sensitive *HNRNPH1* exon 4 skipping isoform. **(I)** Bar plot shows changes in relative expression of *HNRNPH1* in control and ASO-treated cells. Data represented as in (G). **(J)** Bar plot shows changes in MCF7 growth rate upon treatment with ASOs compared to control. Data represented as in (G).

In addition to mutations in individual sites, our experiments in MCF7 cells showcased that also fluctuations in global HNRNPH levels can result in drastic splicing changes. Since HNRNPH has been reported to be dysregulated in several cancers (Chen et al. 2023; Herviou et al. 2020b; Le Bras et al. 2022; Lefave et al. 2011; Li et al. 2016), we wondered whether this is reflected by specific splicing patterns in the tumors. To test this, we analyzed RNA-seq data from the breast invasive carcinoma (BRCA) cohort of The Cancer Genome Atlas (TCGA) (Hutter and Zenklusen 2018). Using the HNRNPH-regulated CE identified in MCF7 cells, we correlated CE inclusion with the expression levels of *HNRNPH1* and *HNRNPH2* in the patient samples. Notably, CE inclusion generally followed *HNRNPH2* expression in the expected direction, with cooperatively enhanced exons tending towards positive correlation whereas cooperatively repressed exons showed predominantly negative correlation (**Figure 7B, Figure S8B, C**). In contrast, we did not detect any correlation with *HNRNPH1* expression as previously reported (Braun et al. 2018) (**Figure 7B**). These findings suggest that the identified CE indeed respond to changes in HNRNPH concentration.

To investigate the clinical relevance of the observed splicing regulation, we used the inclusion frequencies of the HNRNPH-regulated CE for unsupervised clustering of all 1,177 patient samples. Remarkably, the splicing information from these CE was sufficient to clearly distinguish tumor samples (clusters C2–C5) from adjacent normal tissue biopsies (C1; **Figure 7C, D**). Within the tumor samples, the subtypes were sorted by severity, with the most malignant basal subtype showing the most distinct pattern (C5; **Figure 7C, D**). Importantly, the clustering apparently reflected HNRNPH activity: While *HNRNPH1* expression remained relatively stable across clusters, the *HNRNPH2* levels increased from cluster C1 (normal tissue) to C2 (luminal A subtype) and then progressively dropped towards cluster C5 (basal subtype; **Figure 7E**). Compared to randomly picked CE sets, the cooperatively HNRNPH-regulated CE showed a consistently higher performance in separating both normal from tumor samples as well as basal subtype from others (**Figure 7F**). We conclude that variations in HNRNPH concentration modulate splicing-regulatory networks in breast cancer patients. Importantly, *HNRNPH2* expression changes with the breast cancer subtypes, offering HNRNPH-regulated CE as potential biomarkers for subtype differentiation.

### *HNRNPH1* downregulation by antisense oligonucleotides inhibits cancer cell growth

Even though the overall *HNRNPH1* expression remained stable, we observed that skipping of *HNRNPH1* exon 4 showed considerable variation between clusters (**Figure S8D**). Skipping of this exon results in a frameshift and a subsequent premature termination codon which is predicted to target the skipping isoforms to nonsense-mediated mRNA decay (NMD) (Pararajalingam et al. 2020). The relative abundance of the NMD-sensitive isoform was highest in sample cluster C2 and then progressively declined with increasing aggressiveness (C3–C5, **Figure S8D**), while normal tissue samples in cluster C1 showed intermediate levels, suggesting that the availability of functional HNRNPH1 protein is modulated across subtypes. Consistently, an imbalance in *HNRNPH1* isoforms has been linked to inferior outcomes in mantle cell lymphoma (Pararajalingam et al. 2020). Moreover, the *HNRNPH1* NMD isoform followed *HNRNPH2* expression across the five sample clusters, indicating a compensatory cross-regulation between HNRNPH1 and HNRNPH2 in cancer patients.

To confirm that the *HNRNPH1* exon 4 skipping isoform was degraded via NMD in MCF7 cells, we used cycloheximide (CHX) to inhibit translation, resulting in increased *HNRNPH1* levels (**Figure 7G, Figure S8E, F**). To test whether shifting *HNRNPH1* pre-mRNA splicing towards the NMD-sensitive isoform could be used to modulate *HNRNPH1* expression, we designed three different splice-switching antisense oligonucleotides (ASOs) to promote *HNRNPH1* exon 4 skipping and thereby reduce the expression of functional HNRNPH1 (**Figure 7H**). The ASOs were complementary to splicing enhancer sequences surrounding *HNRNPH1* exon 4. Indeed, both ASO1 and ASO2 were sufficient to promote downregulation of *HNRNPH1*, and this effect was revertible under NMD inhibition via CHX (**Figure 7I**). Along with the downregulation of *HNRNPH1*, ASO1 and ASO2 both significantly reduced the proliferative capacity of the MCF7 cells (**Figure 7J**). This suggests that modulating the *HNRNPH1* isoform balance, and hence the availability of functional HNRNPH1 protein, by splice-switching ASOs can interfere with cancer cell growth.

Together, these results demonstrate how cooperative HNRNPH binding, enhanced by rG4 structures, can drive splicing transitions and shape transcriptomic profiles in cancer. They also highlight the potential of targeted splice modulation to alter cell behavior and impact oncogenic splicing programs.

## Discussion

Despite the widespread role of multivalency in protein–RNA interactions, the mechanisms of cooperativity in RNA regulation remain poorly understood. Here, we show that rG4s play a central role in enabling cooperative RNA binding of the splicing regulator HNRNPH, which is critical for the switch-like control of alternative splicing. Using biophysical assays and theoretical modeling, we demonstrate that rG4 unfolding upon HNRNPH binding allows for multivalent, cooperative interactions that translate into highly non-linear splicing responses *in vivo*.

Several previous studies reported that HNRNPH binds to linear G-rich RNAs (Caputi and Zahler 2001; Honoré et al. 1995; Uren et al. 2016). Consistently, HNRNPH has been described to unfold rG4s, to interact with RNA helicases in splicing regulation, and to bind unfolded G-rich RNA (Dardenne et al. 2014; Vo et al. 2022). Moreover, the RNA helicase DHX36 was shown to unwind rG4s to facilitate binding of HNRNPH/F which in turn maintain the rG4 unstructured (Herviou et al. 2020a). Contrasting these results, several studies have suggested that HNRNPH binds to folded rG4 structures (Huang et al. 2017; Neckles et al. 2019). Based on our observations, we propose a unifying model in which HNRNPH recognizes the folded rG4 structures and unfolds these before becoming stably bound to the linear G-rich RNA. rG4 recognition could be mediated by its qRRM domains (Dominguez et al. 2010) or by its IDRs as described e.g. for G3BP1 (He, Yuan, and Wang 2021). Our data thus support a model of a dynamic rG4 folding controlled by RBP binding to modulate RNA regulation.

The role of rG4 structures in mediating indirect cooperativity of HNRNPH binding aligns with emerging evidence that rG4s act as dynamic hubs for RNA regulation (Liu, Chu, and Yang 2022). Our findings extend this paradigm by demonstrating that multiple qRRM domains of HNRNPH cooperatively bind to rG4 structures. This mechanism shares parallels with cooperative RNA binding observed for the proteins FUS and TDP-43, where multi-domain interactions amplify structural remodeling (Huang et al. 2017). Our biophysical model provides a route for understanding how spatial coordination between qRRMs enhances binding avidity, consistent with structural studies of multi-modular RNA-binding. This mechanism may explain how HNRNPH achieves sequence-specific recognition despite its broad RNA binding preference. So far, only few examples describe how indirect cooperativity can impact RNA binding (Becker et al. 2019; HafezQorani et al. 2016). Our data suggest rG4-mediated cooperativity in HNRNPH regulation occurs on a transcriptome-wide scale, establishing indirect cooperativity as a global phenomenon paralleling direct cooperative mechanisms such as multivalent protein–protein interactions (Daugherty et al. 2010; Gueroussov et al. 2017; Rengifo-Gonzalez et al. 2021).

The link between molecular binding dynamics and switch-like splicing regulation underscores a fundamental principle of gene expression: kinetic cascades amplify subtle binding differences into robust cellular decisions. Our kinetic model of splicing decision-making shows how the sequential cooperative steps—from RNA binding to spliceosome assembly to exon inclusion—converge into a signal amplification cascade and suggests that combinatorial cooperativity and delayed feedback loops could explain the “digital” splicing patterns observed in cancer cells (David and Manley 2010).This mirrors mechanisms in signal transduction and transcriptional networks (Bintu et al. 2005) but has been understudied in splicing contexts. Our framework explains how modest cooperativity at each step can yield the steep dose–response curves observed for HNRNPH-regulated exons. This aligns with reports on other splicing factors, for which cooperative RNA regulation has been observed, such as MBNL1 (Wagner et al. 2016). It may further extend to other RNA-regulatory processes that require stepwise complex assemblies, such as cleavage and polyadenylation.

An interesting question arising is what might be the benefits of cooperativity in splicing regulation? In general, cooperativity allows for sharper, more switch-like responses to regulatory signals in gene regulation (Gutierrez, Monteoliva, and Diambra 2012). Changes in expression are buffered until a certain threshold, but going beyond that threshold leads to a pronounced effect. Such switch-like regulation is for example implemented during gene regulatory processes in early Drosophila development and enables sharp boundaries of gene expression within the embryo (Lopes, Spirov, and Bisch 2012). Furthermore, cooperativity can promote synchronized cellular responses and can prevent crosstalk between gene regulatory pathways (Friedlander et al. 2016). Recently this principle could also be used in synthetic gene regulatory networks to reduce “off-target” regulation (Bragdon et al. 2023). In the case of HNRNPH, cooperativity could contribute to control important regulatory decisions for example in the context of neuronal differentiation (Grammatikakis et al. 2016), or apoptosis and cell motility (Lefave et al. 2011).

Importantly, HNRNPH expression and function have been implicated in cancer progression (Chen et al. 2023; Herviou et al. 2020a; Le Bras et al. 2022; Lefave et al. 2011; Li et al. 2016). However, unlike many oncogenes, dysregulation of RBP expression levels appears relatively modest across various cancer types (Wang, Liu, and Shyr 2015). For HNRNPH this discrepancy might be explained by the cooperative, switch-like splicing regulation mediated by HNRNPH. Supporting this notion, we observed that relatively subtle variations in HNRNPH expression across breast cancer subtypes are associated with coordinated splicing changes. These changes not only distinguish between tumor subtypes but also underscore the critical role of HNRNPH in shaping cancer-specific splicing landscapes, generalizing previous findings on MCL-1 and HER2 splicing in breast cancer (Gautrey et al. 2015; Tyson-Capper and Gautrey 2018).

Reducing HNRNPH expression levels has been shown to impair the viability of breast cancer cells (Tyson-Capper and Gautrey 2018). Our findings suggest that ASOs promoting non-productive *HNRNPH1* splicing could offer similar therapeutic benefits, highlighting the potential of splice-switching ASOs to target dysregulated HNRNPH expression in cancer. Our findings extend to the clinical level, demonstrating that HNRNPH-dependent splicing events can serve as diagnostic markers to events stratify breast cancer subtypes. Together, these insights open new avenues for investigating HNRNPH-targeted therapies and underscore the clinical relevance of splicing regulation in cancer.

Altogether, our findings establish rG4-mediated cooperativity as a mechanistic principle for modulating RNA–protein interactions and splicing outcomes. The observations highlight how RNA secondary structures and kinetic integration across regulatory layers can generate ultrasensitive control of gene expression, with implications for both biomarker discovery and targeted cancer therapies.

## Methods

### Cell culture and transfection of plasmids and siRNAs

MCF7 cells were purchased from ATCC and cultured in RPMI (Life Technologies) supplemented with 10% fetal bovine serum (Life Technologies), 1% L-Glutamine (Life Technologies), and 10 µg/ml insulin (Sigma-Aldrich). The cells were kept at 37 °C with 5% CO_2_ and regularly tested for mycoplasma infection.

For transfection of plasmids and siRNAs, cells were seeded one day prior to transfection with 5*10^5^ cells (plasmid transfection) or 4*10^5^ cells (siRNA transfection) per well in a 6-well plate. To achieve gradual HNRNPH1 overexpression, 0.6 µg, 1 µg or 2.5 µg pcDNA3.1 (+) HNRNPH1 overexpression construct was transfected. Plasmid transfection was conducted using 5 µL Lipofectamine 2000 (Invitrogen) in 200 µk OPTI-MEM (Invitrogen) and 20 min incubation before addition to the cells. As a control, the empty pcDNA3.1 (+) vector was transfected. For *HNRNPH1* KD, single small interfering RNA (siRNA) with a final concentration of 20 nM was used (**Supplementary Table S6**) (Rauch et al. 2011). A non-targeting siRNA was transfected as control (**Supplementary Table S6**). siRNA transfection was performed with 5 µL RNAiMax (Invitrogen) in 200 µl OPTI-MEM following the same procedure as for plasmid transfection. Cells were harvested 48 h after plasmid or siRNA transfection. For siRNA titration experiments, increasing concentrations of the HNRNPH1-targeting siRNA were used.

In the case of splicing minigene experiments, 4*10^5^ cells per well were seeded in a 6-well plate and transfected 24 h later with HNRNPH1-targeting siRNA. Another 24 h later, the cells were transfected with 2 µg minigene plasmid using Lipofectamine 2000 as described above. The cells were harvested 72 h after seeding.

### Cloning

To clone the *AKR1A1* minigene construct, the genomic region of interest was PCR-amplified from human genomic DNA using the Phusion DNA polymerase (New England Biolabs). The primers were extended with either a *Hind*III (forward primer) or an *Xba*I (reverse primer) restriction enzyme site (**Supplementary Table S6**). The ∼1,100 bp long PCR product was gel-purified with the QIAquick Gel Extraction Kit (QIAGEN). The PCR product as well as the donor vector pcDNA 3.1 (+) were digested with the *Hind*III-HF and *Xba*I restriction endonucleases (New England Biolabs), 20 units (U) each, and afterward again gel-purified. The ligation reaction was performed using the T4 DNA Ligase (New England Biolabs) following the manufacturer’s instruction with a 1:3 vector:insert ratio. To achieve a minigene splicing pattern that was comparable to the endogenous *AKR1A1* splicing, the 3’ splice site of the minigene was strengthened by the introduction of one point mutation (G549T) using the Q5 Site-Directed Mutagenesis Kit (New England Biolabs) according to the manufacturer’s recommendations (**Supplementary Table S6**). From our previous work, we used the pcDNA3.1 (+) HNRNPH1 overexpression construct (Braun et al. 2018).

The HNRNPH1 expression construct for recombinant protein production was generated using Gibson assembly. All primers for Gibson assembly reactions were designed with the NEBuilder Assembly tool (New England Biolabs). Gibson reactions were set up by mixing 100 ng DNA (vector:insert ratio 1:3) with 10 µl of Gibson assembly 2x master mix from the IMB Protein Production Core Facility. The Gibson reactions were incubated at 50 °C for 1 h and afterward transformed in Dh5α *E. coli*. To generate the MBP-HNRNPH1-His_6_ expression construct, the HNRNPH1 open reading frame (ORF) was cloned into a pET vector backbone carrying a C-terminal His_6_-tag and an N-terminal MBP-tag, a kind gift from the IMB Protein Production Core Facility. The HNRNPH1 ORF was PCR-amplified from our pcDNA3.1 (+) HNRNPH1 overexpression construct using PCR (**Supplementary Table S6**). The pET vector backbone was prepared for Gibson assembly performing a PCR reaction (**Supplementary Table S6**).

### *AKR1A1* minigene mutagenesis

For the splicing analysis of the *AKR1A1* minigene, two HNRNPH binding sites (chr1:45,568,349-45,568,431:+ and chr1:45,568,702-45,568,761:+) in the upstream and downstream intron were mutated (**Supplementary Table S7**). Therefore, for each triple G-run (GGG) the middle G was substituted and for each quadruple G-run (GGGG) the two middle G were substituted. The mutations were performed using the Q5 Site-Directed Mutagenesis Kit (New England Biolabs) according to the manufacturer’s recommendations (PCR oligos available in **Supplementary Table S6**). After mutation of the endogenous HNRNPH binding sites, we generated two new *AKR1A1* minigene constructs. We introduced two new HNRNPH binding sites in the downstream intron of the *AKR1A1* minigene (**Supplementary Table S7**). The two new HNRNPH binding sites were chosen from our RT-Stop profiling data and did not show any rG4 formation (Non-G4 #1: CTGGGGTGATTTGGGCAGTCTCCGGG, Non-G4 #2: CGGGTGGTTGGTT). To introduce the new binding sites, we used the Q5 Site-Directed Mutagenesis Kit (New England Biolabs) (PCR oligonucleotides available in **Supplementary Table S6**).

### Semiquantitative RT-PCR for splice isoform quantification

Semiquantitative RT-PCR was conducted to quantify splicing patterns of endogenous mRNA or minigene splicing events. Total RNA was isolated from cells with the RNeasy Plus Mini Kit (Qiagen) following the manufacturer’s instructions. Then, the reverse transcription was conducted using 500 ng RNA with RevertAid Reverse Transcriptase (Thermo Fisher Scientific) and Oligo(dT)_18_ primer (Thermo Fisher Scientific) following the manufacturer’s instructions.

To quantify splicing patterns of *AKR1A1* splicing events, 1 µl of the cDNA was used as input for the PCR reaction with OneTaq DNA polymerase (New England Biolabs) (PCR conditions: 94 °C for 30 s, 30 cycles [endogenous] or 25 cycles [minigene] of [94 °C for 20 s, 55 °C for 30 s, 68 °C for 30 s] and final extension at 68 °C for 5 min). To amplify the *AKR1A1* splicing isoforms the forward primer was either combined with the endogenously binding reverse primer or with the reverse primer binding the pcDNA3.1 (+) backbone upstream of the polyadenylation site (**Supplementary Table S6**).

To validate splicing events from the RNA-seq analysis of the gradual downregulation and gradual overexpression of HNRNPH in MCF7 cells, RT-PCR was performed for *MYL6*, *PIP4K2C* and *SNHG21* (gradual downregulation from (Braun et al. 2018)). Therefore, 1 µL of cDNA was used as input for a 30-cycle PCR reaction with OneTaq DNA polymerase (New England Biolabs) (PCR conditions described above using adapted annealing temperatures; **Supplementary Table S6**).

The TapeStation 2200/4400 capillary gel electrophoresis instrument (Agilent) was used for isoform quantification of the PCR products on D1000 tapes.

### Quantitative real-time PCR (qPCR)

For quantitative determination of mRNA levels, 500 ng total RNA was reversely transcribed into cDNA using the RevertAid Reverse Transcriptase (Thermo Fisher Scientific) and Oligo(dT)_18_ primer (Thermo Fisher Scientific) following the manufacturer’s instructions. In accordance to the manufacturer’s instruction, qPCR reactions were performed in technical triplicates using the Luminaris HiGreen qPCR Master Mix, low ROX (Thermo Fisher Scientific) with forward and reverse primer (0.3 µM each; **Supplementary Table S6**) and 2 µl of 1:10 diluted cDNA as template. All qPCR reactions were run on a ViiA 7 Real-Time PCR System (Applied Biosystems). For normalization, mRNA levels of the housekeeping gene *beta*-*actin* were determined (**Supplementary Table S6**). The quantification of mRNA expression levels was performed with the ΔΔCt method (Livak and Schmittgen 2001).

### Oligonucleotide library for *in vitro* transcripts (IVT library)

The IVT library consists of 200-nt oligonucleotide sequences composed of an 18-nt T7 promoter, a 146-nt transcript region around an HNRNPH binding site and/or predicted RNA G-quadruplex (rG4), a 15-nt unique barcode and a 21-nt L3 linker sequence for sequencing. Barcodes were chosen to have a Hamming distance ≥ 5 to any other barcode and contain no more than two consecutive Gs. The chosen transcript regions are located in cooperatively regulated or control splicing events that had been identified in preliminary analyses. Briefly, HNRNPH binding sites and predicted rG4s (G-tract length > 2) were determined and merged into a single region in the case of overlaps, resulting in 1,666 regions (each composed of a binding site and/or rG4) from regulated splicing events complemented by 1,334 regions from control events. The 3,000 regions and their up- and downstream flanking transcript regions (total length of 146 nt) were placed downstream of the T7 promoter. The flanking transcript regions were chosen such that the sites of interest (binding site or rG4) always start at position 51 in the 200-nt sequence. Each of the 3,000 regions is present four times in the IVT library with different barcodes, resulting in 12,000 oligonucleotides. Note that 106 regions (with 4 barcodes each) were excluded from the subsequent analysis, resulting in a total of 11,576 oligonucleotides that were processed further.

The single-stranded (ss)DNA oligonucleotide library was ordered from TWIST Bioscience as an oligonucleotide pool, Tier 3L ssDNA with 12,000 individual oligonucleotides of 200 nt.

### PCR amplification of ssDNA oligonucleotides

The sequences of interest were ordered as ssDNA oligonucleotides from Integrated DNA Technology (IDT) or as an ssDNA oligonucleotide pool from TWIST Bioscience (**Supplementary Table S4**). All oligonucleotides contained the T7 promotor sequence at their 5’ end. The ssDNA was PCR-amplified with Phusion DNA polymerase to have double-stranded (ds)DNA for *in vitro* transcription (PCR conditions: 98 °C for 30 s, 25 cycles [individual ssDNA oligonucleotides] or 10 cycles [ssDNA oligonucleotide pool] of [98 °C for 30 s, 50 °C for 30 sec, 72 °C for 20 s] and final extension at 72°C for 10 min) (**Supplementary Table S6**). The PCR reactions were set up with 25 ng ssDNA oligonucleotide or 10 ng ssDNA oligonucleotide pool, 1 U/50 µl Phusion DNA polymerase (New England Biolabs), 1x Phusion GC buffer (New England Biolabs), 200 µM dNTPs (New England Biolabs), 0.5 µM forward primer and 0.5 µM reverse primer in total 50 µl (ssDNA oligonucleotide) or 200 µl (ssDNA oligonucleotide pool) reaction volume.

### MyONE Silane bead clean-up

After PCR amplification, the dsDNA was purified using MyONE Silane beads (Invitrogen). Briefly, 20 µl MyONE Silane beads per sample were magnetically separated and washed with 500 µL RLT buffer (Qiagen). After washing, the beads were again magnetically separated, and the supernatant was removed. The beads were resuspended in 250 µl RLT buffer per sample, added to the PCR reaction and mixed. Then, 1 volume of 100% ethanol (50 µl PCR product + 250 µl beads : 300 µl 100% ethanol) was added, carefully mixed, and incubated for 5 min at room temperature (RT). After incubation, the mixture was again mixed and incubated for further 5 min. To remove the supernatant, the beads were magnetically separated and washed with 1 ml of 80% ethanol. After transferring the samples to a new tube, the beads were washed twice in 80% ethanol. Finally, the beads were magnetically separated, and the supernatant was removed completely. The beads were air-dried for 5 min at RT. Then, the beads were resuspended in 12 µl nuclease-free H_2_O and incubated for 5 min at RT. The beads were magnetically separated, and the eluted dsDNA was transferred to a new tube for *in vitro* transcription. Eluted DNA was quality-controlled using the TapeStation 2200 capillary gel electrophoresis instrument (Agilent) on D1000 tapes.

### *In vitro* transcription

For *in vitro* transcription, the HiScribe T7 High Yield RNA Synthesis Kit (New England Biolabs) was used following the manufacturer’s recommendation for short transcripts (<0.3 kb) in a total volume of 40 µl with 500-1000 ng of template dsDNA. To generate RNA with 7-deaza-GTP instead of GTP, 7.5 mM GTP was substituted with 6.75 mM 7-deaza-GTP (TriLink Biotechnologies) and 0.75 mM GTP. All *in vitro* transcription reactions were incubated in a thermocycler at 37 °C for 16 h. To remove template DNA, the *in vitro* transcription reactions were treated with 4 U Turbo DNase (Invitrogen) in 100 µl reaction volume for 15 min at 37 °C. Subsequently, the RNA was purified using the RNeasy Plus Mini Kit (Qiagen) following the manufacturer’s instructions for the purification of total RNA containing miRNA. The RNA was eluted in 20 µl nuclease-free water and the length was validated using the TapeStation 2200 system with RNA ScreenTapes.

### Western blot

For protein level quantification, cells were harvested and lysed in modified RIPA buffer (50 mM Tris-HCl pH 7.5, 150 mM NaCl, 1 mM EDTA, 1% NP-40, 0.1% sodium deoxycholate, protease inhibitor cocktail; Roche). The protein lysates were run on NuPAGE 4-12% Bis-Tris gels (Invitrogen). For Western blot analysis, the following antibodies were used: rabbit polyclonal anti-HNRNPH, 1:10,000 dilution (AB10374, Abcam) and mouse monoclonal anti-Vinculin, 1:10,000 dilution (V9624, Sigma-Aldrich).

### iCLIP experiments

iCLIP2 was performed to gain information about the RNA binding profile of HNRNPH in MCF7 cells following the published protocol (Buchbender et al. 2020). Briefly, the iCLIP libraries were made from MCF7 cells with approximately 1.5*10^6^ cells per immunoprecipitation in seven replicates. To pull down HNRNPH from the cell lysates, 5 µg of a polyclonal rabbit anti-HNRNPH antibody from Abcam (AB10374) was used. RNase digestion was performed with a 1/250 diluted RNase I (Ambion). For reverse transcription, the primer 5’-GGATCCTGAACCGCT-3’ was used. We sequenced the iCLIP library on an Illumina NextSeq 500 platform with high output and 75 cycles.

### RTstop profiling

To map the positions of RNA G-quadruplex (rG4) structures *in vitro* in high throughput, the RTstop profiling protocol was adapted (Guo and Bartel 2016). 1 µg RNA of the IVT library was used as a template for the reverse transcription with SuperScript III (Invitrogen). First, the RNA was mixed with 0.5 pmol of the RT primer (5’-GGATCCTGAACCGCT-3’) and 0.5 mM dNTPs in reverse transcription buffer (20 mM Tris-HCl pH 7.5, 150 mM KCl or 150 mM NaCl, 3 mM MgCl_2_) in a total reaction volume of 19.5 µl. The mixture was incubated at 80 °C for 2 min, followed by incubation at 25 °C for 2 min. Then, 100 U of SuperScript III enzyme was added, and the reaction was incubated for 10 min at 42 °C. To hydrolyze the template RNA, 1.65 µl of 1 M NaOH solution was added, followed by incubation at 98 °C for 20 min. The addition of 20 µl 1 M HEPES-NaOH pH 7.3 stopped the hydrolysis reaction. From here on, the library preparation of the iCLIP2 protocol was used continuing with the second adapter ligation to the 3’ end of the cDNA (Buchbender et al. 2020). In brief, after MyONE Silane bead clean-up, the second adaptor was ligated to the 3’ end of the cDNA on the beads. Then, the cDNA was purified via MyONE Silane beads and the library was amplified using PCR (**Supplementary Table S6**). After two rounds of ProNex size selection with a sample-to-ProNex (v/v) ratio of 1:2.4, the final library was sequenced on an Illumina NextSeq 500 sequencing machine as 141-nt + 25-nt paired-end reads. The first read (R1) includes a 6-nt sample barcode as well as 5+4-nt unique molecular identifiers (UMIs) at the beginning, while the barcode of the oligonucleotides is found at the first 15-nt of the second read (R2).

### Recombinant protein expression

Recombinant His_6_-MBP-tagged, full-length HNRNPH1 was expressed in *E. coli* BL21 (DE3) pLysS using a pET vector. Cells were grown at 37 °C and 160 rpm until an OD_600_ of 0.6. Subsequently, recombinant protein expression was induced by addition of IPTG (1 mM final concentration). The expression culture was further incubated at 37 °C and 160 rpm for 3 h. Cells were harvested by centrifugation at 4000 xg for 15 min at 4 °C and resuspended in lysis buffer (50 mM Tris-HCl pH 7.5, 500 mM NaCl, 5% glycerol, 10 mM imidazole) supplemented with 0.1% Triton X-100, protease inhibitor, Sm nuclease (6 cU/µl), 1 mM DTT and 1 mM MgCl_2_. Cells were lysed by sonication, the lysate was cleared by centrifugation (45,000 xg, 30 min, 4 °C) and then passed over a HisTrap 5 mL column (GE Healthcare) to retain the recombinant protein. After washing the column with lysis buffer, the bound recombinant protein was eluted using a gradient from 10 to 500 mM imidazole in lysis buffer over 10 column volumes. The HNRNPH1-containing elution fractions were pooled and run on a HiLoad 16/60 pg Superdex 200 column (GE Healthcare) in gel filtration buffer (30 mM Na-HEPES pH 7.4, 150 mM NaCl,1 mM DTT, 5% glycerol). The fractions containing the recombinant protein were pooled and snap-frozen in liquid nitrogen for further storage at –80 °C.

### 5’ dephosphorylation of RNA

For radioactive labelling, RNA was dephosphorylated at the 5’ end. Therefore, the RNA was incubated with 0.5 µl RNasin Plus (Promega) and 25 U antarctic phosphatase (New England Biolabs) in 20 µl for 30 min at 37 °C. Subsequently, the RNA was purified using the RNA Clean & Concentrator-5 Kit (Zymo Research) following the manufacturer’s instructions and aiming for a final RNA concentration of 15 µM.

### In vitro binding

To analyze the binding conditions of HNRNPH1 to RNA, the RNA was first radioactively labelled. 5 µM of 5’-dephosphorylated RNA was incubated with 1 µl ^32^P-γ-ATP, 10 U T4 Polynucleotide Kinase (New England Biolabs) and 0.5 µl RNasin Plus (Promega) in a total volume of 20 µl for 20 min at 37 °C and 1100 rpm. Then, the radioactively labelled RNA was purified using Sephadex G-25 quick spin columns (Roche) following the manufacturer’s instructions. To re-fold the RNA, 100 nM labelled RNA was first incubated with 150 mM KCl or 150 mM NaCl and 20 mM Tris-HCl pH 7.5 in total 13 µl for 5 min at 80 °C followed by 15 min at 25 °C. Subsequently, 2 µl 10x protein binding buffer (100 mM HEPES pH 7.2, 30 mM MgCl_2_, 30% glycerol, 10 mM DTT) and 5 µl 4x concentrated recombinant HNRNPH1 protein were added. The RNA-protein reaction was incubated at 37 °C for 10 min and 1,100 rpm. Then, the reaction was irradiated with 250 mJ/cm^2^ in a UV-C crosslinker (CL-1000 Ultraviolet Crosslinker, UVP) at 254 nm. The samples were run on NuPAGE 3-8% Tris-Acetate gels (Invitrogen) for 60 min at 150 V and afterwards the protein-RNA complexes were transferred to a nitrocellulose membrane for 60 min at 30 V. Finally, the membrane was exposed to a BAS Storage Phosphor Screen (GE Healthcare) for 2 h and then scanned on a Typhoon scanner (GE Healthcare).

The *in vitro* binding of recombinant HNRNPH1 protein to RNA was analyzed in high throughput using *in vitro* iCLIP (Buchbender et al. 2020). Therefore, we crosslinked recombinant HNRNPH1 protein to the IVT library and performed iCLIP library preparation. In brief, 100 nM radioactively labelled RNA was re-folded in 150 mM KCl and 20 mM Tris-HCl pH 7.5 for 5 min at 80 °C followed by 15 min at 25 °C. Then, the RNA was incubated with 500 nM recombinant HNRNPH1 protein for the *in vitro* binding. Following the *in vitro* binding, the library preparation for sequencing was performed according to the previously published iCLIP protocol, continuing with the RNA isolation from the nitrocellulose membrane (Buchbender et al. 2020). To normalize the final *in vitro* binding signal, each sample included 0.1 nM of a spike-in pool (10 IVTs) that was crosslinked with 500 nM recombinant HNRNPH1 protein. The experiment was done in four replicates for the IVT library with GTP *in vitro* transcription and for the IVT library with 7-deaza-GTP *in vitro* transcription. The final library was sequenced with on an Illumina NextSeq 500 sequencing machine as 141-nt + 25-nt paired-end reads. The first read (R1) includes a 6-nt sample barcode as well as 5+4-nt unique molecular identifiers (UMIs) at the beginning, while the barcode of the oligonucleotides is found at the first 15-nt of the second read (R2).

### Circular dichroism (CD) spectroscopy

All CD spectra were measured with a Jasco J-1500 CD spectrometer (Jasco). The RNA oligonucleotides were ordered from Integrated DNA Technologies (IDT) and dissolved in nuclease-free water to a final concentration of 100 µM (**Supplementary Table S6**). Prior to the spectroscopic analysis, 10 µM RNA in 20 mM Tris-HCl pH 7.5 in a total volume of 200 µl was heated to 95 °C for 3 min in the presence of either 150 mM KCl, 150 mM NaCl or without any monovalent cation, and then gradually cooled to RT for 1 h. CD spectra of these samples were scanned at 25 °C at a scan rate of 200 nm/min, with a 1 sec response time, 1 nm bandwidth and in continuous scan mode in 1 mm cuvettes. The CD spectra presented are averages of 6 individual spectra corrected for buffer background.

### Fluorescence resonance energy transfer (FRET)

All fluorescence measurements were performed with a Tecan Infinite M200 Pro (Tecan). The FRET-RNA oligonucleotides were ordered from IDT and dissolved in nuclease-free water (**Supplementary Table S6**). To re-fold the RNA, 0.44 µM RNA in 20 mM Tris-HCl pH 7.5 was heated at 95 °C for 3 min and gradually cooled to RT for 1 h in the presence of either 150 mM KCl or 150 mM LiCl. 0.4 µM re-folded RNA was incubated with 100 nM, 250 nM or 500 nM recombinant HNRNPH1 protein in a total volume of 50 µl for 10 min at room temperature. The RNA–protein mixture was transferred into a white 96-well plate (Thermo Fisher Scientific) and fluorescence was measured from top. Excitation (bandwidth: 9 nm) was performed with 25 flashes at 485 nm and emission (bandwidth: 20 nm) was detected at 520 nm. Each well was measured four times using an optimal gain. All measurements were performed in five independent replicates.

### RNA-seq data processing

RNA-seq libraries were sequenced on an Illumina NextSeq 500 sequencing machine as 159-nt single-end reads. Basic quality controls were done for all RNA-seq samples with FastQC (version 0.11.5) (https://www.bioinformatics.babraham.ac.uk/projects/fastqc/). Prior to mapping, all reads were trimmed to 100-nt using basic Bash commands. Trimmed reads were mapped using STAR (version 2.5.4b) (Dobin et al. 2013) allowing up to 5 mismatches (--outFilterMismatchNmax 5) and a splice junction overhang (--sjdbOverhang) of 99-nt. Genome assembly and annotation of GENCODE (Dobin et al. 2013) release 25 were used during mapping. Subsequently, secondary hits were removed using Samtools (version 1.5) (Danecek et al. 2021). Uniquely mapped exonic reads per gene were counted using featureCounts of the Subread tool suite (version 1.5.1) (Liao, Smyth, and Shi 2014) with non-default parameters --donotsort -s2.

### Expression analysis

Genes differentially expressed between the groups were detected using the R/Bioconductor package DESeq2 (version 1.22.2) (Love, Huber, and Anders 2014) in an R environment (version 3.5.1) (R Core Team 2020).

### Alternative splicing analysis

Identification and quantification of alternative splicing events was performed using the MAJIQ and VOILA packages (both version 2.3).(Vaquero-Garcia et al. 2023) First, *majiq build* was used to generate a splice graph for each gene and to detect so-called local splicing variations (LSVs) in the splice graphs by taking the BAM files of all titration experiments (except replicate 3 of KD1000 and KD5000) and the human GENCODE gene annotation (release 39) as input. In the second step, changes in the splice junction usage and retention of introns (ΔPSI) were quantified between the knockdown and matched control experiment as well as between the overexpression and matched control experiments using *majiq deltapsi*. In the final step, *voila modulize* was used to reconstruct binary alternative splicing events (e.g., cassette exon or intron retention) from the LSVs and to calculate the probability that the |ΔPSI| of a junction or intron exceeds a value of 0.02 (--changing-between-group-dpsi 0.02).

In total, *voila modulize* reports 14 different types of binary alternative splicing events reconstructed from either one (e.g., intron retention) or two LSVs (e.g., cassette exon), with two junctions reported for each LSV. An alternative splicing event is considered significantly regulated in a pairwise comparison if: (i) the two junctions of one LSV have a probability of 90% that the change was 0.02 or more (P(|ΔPSI| ≥ 0.02) ≥ 0.9), (ii) the junctions of all LSVs have a change of at least 0.025 (|ΔPSI| ≥ 0.025), (iii) junctions of the same LSV show opposite regulation and (iv) the less regulated junction of a LSV shows at least 50% of the change of the more regulated junction. The exception of this procedure are multi-exon skipping events, where rule (iii) cannot be applied as the two junctions of each LSV have the same reported ΔPSI value.

Downstream analyses focused on cassette exon, intron retention and alternative last exon events. We considered events of these three binary event types that had ΔPSI values in all comparisons and that were significantly regulated in at least one of the two strongest knockdown or overexpression experiments. For each of the events, we determined a representative junction whose ΔPSI values were used to describe the inclusion levels of the alternative exon, intron or first alternative last exon. For CE events, we used the inclusion junction (constitute exon 1 or to alternative exon, C1_A or, C2_A, respectively) that showed the strongest change in the two strongest knockdown and overexpression experiments, whereas for intron retention events it was the intron itself (C1_C2_intron or C2_C1_intron) and for the alternative last exon events the proximal junction (Proximal). In addition, a set of control cassette exon events was determined in which no junction had a change greater than 2.5% (|ΔPSI| | ≤ 0.025). Here, the first inclusion junction (C1_A) was selected as a representative.

### Dose–response curves

The ΔPSI values of the cassette exon, intron retention and alternative last exon events were used together with the relative HNRNPH protein levels to fit a four parametric logistic curve (LL.4(names = c(“n_H_”, “min”, “max”, “ec50”)) using the drm() function of the drc (version 3.0-1) R package (Ritz et al. 2015). The following constraints were used for parameter ranges: n_H_ ∈ [–Inf, +Inf], min ∈ [–max(PSI_KD_Control_, PSI_OE_Control_), 0], max ∈ [0, 1-min(PSI_KD_Control_, PSI_OE_Control_)] and ec50 ∈ [0, +Inf]. As a goodness of fit the pseudo R^2^ was calculated using the R2nls function of the R package aomisc (version 0.674) (https://github.com/OnofriAndreaPG/aomisc). Events with a pseudo R^2^ ≥ 0.75 were subsequently classified into four categories based on the degree of cooperativity quantified by the estimated Hill coefficient (n_H_). The four categories included: (i,ii) cooperatively regulated splicing events (*n*_*H*_ ≥ 2) that were (i) enhanced or (ii) repressed by HNRNPH, and (iii,iv) non-cooperatively regulated events (*n*_*H*_ < 2) that were (iii) enhanced or (iv) repressed by HNRNPH.

### iCLIP data processing

Basic quality controls were done with FastQC (version 0.11.5) (https://www.bioinformatics.babraham.ac.uk/projects/fastqc/) and reads were filtered based on sequencing qualities (Phred score) in the sample barcode and UMI regions. All reads with two or more positions with a Phred score below 20 in the sample barcode or a position with a Phred score below 17 in the UMIs were removed from further analysis. Reads were de-multiplexed based on the sample barcode, which is found on positions 6 to 11 of the reads, using Flexbar (version 3.0.0) (Roehr, Dieterich, and Reinert 2017). Subsequently, adapter sequences were trimmed from read ends using Flexbar requiring a minimal overlap of 1-nt of read and adapter. Barcode and UMI sequences were also removed from the reads and added to the read identifiers. Trimmed reads shorter than 15-nt were removed from further analysis.

Remaining reads were mapped to the human genome (hg38/GRCh38.p7) and its annotation (GENCODE release 25) (Frankish et al. 2019) using STAR (version 2.5.4b) (Dobin et al. 2013). When running STAR, up to two mismatches were allowed, soft-clipping was prohibited on the 5’ end of the reads and only uniquely mapped reads were kept for further analysis. Duplicate reads were marked using the dedup function of bamUtil (version 1.0.13) (Jun et al. 2015), which defines duplicates as reads whose 5’ ends map to the same position in the genome. Using an in-house Perl script, we then removed all marked duplicates with identical UMIs representing technical duplicates, while biological duplicates with different UMIs were kept.

The nucleotide position upstream of aligned reads is considered as the ‘crosslink nucleotide’, with each read counted as individual ‘crosslink event’. Duplicate removed BAM files were transformed to BED files using bamToBed of the BEDTools suite (version 2.25.0) (Quinlan and Hall 2010), which were then reduced to just the position upstream of the reads using basic Bash commands. BedGraph files were generated by counting reads per position. They were split by strand and further transformed to bigWig file format using bedGraphToBigWig of the UCSC tool suite kentUtils (version 365) (https://github.com/ENCODE-DCC/kentUtils/).

### Binding site definition

The BAM files of the seven independent iCLIP replicates were combined into two replicates by merging 3 and 4 of the files, respectively, into a single BAM file each using Samtools (version 1.9) (Li et al. 2009). The two replicate BAM files served as input for the subsequent peak calling using PureCLIP (version 1.3.1)(Krakau et al. 2017) with default parameters.

Raw HNRNPH crosslink events from the 7 iCLIP experiments were loaded from bigWig files and then merged by independently summing the number of crosslink events per genomic position for the plus and minus strands. The merged raw crosslink events were used together with the significant crosslink positions reported by PureCLIP to iteratively define 5-nt wide HNRNPH binding sites. As a first step, significant crosslink positions with a distance of 3 nt or less (≤ 3 nt between the sites) were merged into regions potentially containing the binding sites, and regions shorter than 2 nt immediately removed. Next, a two-strep procedure was applied to iteratively place 5 nt wide HNRNPH binding sites into the regions: To place the first binding site into a region, the position with the highest number of crosslink events was determined for each region, extended by 2 nt up- and downstream and stored as 5 nt binding site. Then, the 5 nt of the binding sites along with additional 4 nt on either side were cut out from the regions, leaving behind up to two remaining regions on either side. These two steps were repeated until no more binding sites could be placed. In a final cleanup step, binding sites whose center was not a significant crosslink position or the position with the highest number of crosslink events were removed, as well as binding sites with less than 3 positions with crosslink events.

### Genomic distribution of HNRNPH binding sites

Annotation obtained from GENCODE (release 39) was filtered for the gene support (≤ 2) and transcript support level (≤3 and NA). The binding sites were then overlapped with all remaining genes to assign a biotype to each binding site. In the case of overlap with multiple genes of different biotypes, the hierarchy protein-coding > lncRNA > other was used. For binding sites that overlapped protein-coding genes, we further assigned the overlapping transcript region. For multiple overlapping regions, we assigned the one that occurs in more isoforms, and in the case of a tie, we used the hierarchy CDS > 5’ UTR > 3’ UTR > intron.

To further elucidate the dependence between the expression of a gene and binding by HNRNPH (at least one binding site on the gene), we determined the proportion of genes bound and the number of genes expressed at different TPM cutoffs (0 to 80 in 1 TPM increments). In this context, we also determined the proportion of expressed genes (TPM ≥ 1) bound by HNRNPH for each biotype.

In a similar way we overlapped genes with at least one significant alternative splicing event and genes bound by HNRNPH by applying an expression cutoff of TPM > 1. We tested the overlap of genes to be significant with Fisher’s exact test. For the cassette exon events, we further determined the number of overlapping HNRNPH binding sites for the cooperatively enhanced, cooperatively repressed and control events. We considered the genomic location of the events to be the region from the first position of the upstream exon to the last position of the downstream exon.

### HNRNPH RNA splicing maps on cooperatively regulated events

To identify regions of enriched binding in the cooperatively enhanced and repressed events using RNA splice maps, we first picked two sets of control events that matched the number of enhanced and repressed events, respectively. An important aspect in the selection of control events is the preservation of the PSI distribution, which solves the problem that exons with differences in inclusion levels but bound with similar affinity show strong differences in the number of measured crosslink events. Therefore, for both the cooperatively enhanced and repressed events, we sampled two independent sets of PSI-matched control events. First, five quantiles of the PSI distributions were determined (0, 5, 70, 80 and 100% for enhanced and 0, 10, 65, 90 and 100% for repressed events). The PSI values obtained for the quantiles served as boundaries for four PSI intervals to which the control events were assigned based on their PSI values. In the last step, a fixed number of control events was sampled from each PSI interval, such that the resulting sets of control events approximately had the same size and PSI distribution as the enhanced and repressed events, respectively.

RNA splicing maps were constructed around the 5’ splice site of the upstream exon, the 3’ and 5’ splice sites of the alternative exon and the 3’ splice site of the downstream exon. The first step was to open a window up to 50-nt into exons and up to 300-nt into introns at each splice site, determine the number of crosslink events at each window position and normalize the number of crosslink events to a range of 0 to 1 (min–max normalization) to counteract the effect of few windows with very high numbers of crosslink events. For the final RNA splicing maps, the normalized crosslink events at each window position were averaged over all exons, followed by loess smoothing. The latter two steps were performed separately for the cooperatively enhanced, cooperatively repressed and two sets of PSI-matched control exons.

Signal differences between the cooperatively enhanced or repressed events and their PSI-matched control events were determined in overlapping 10-nt windows (step size of 1). In brief, in each 10-nt window the average number of normalized crosslink events was determined for the individual cooperatively regulated and PSI-matched control events. The resulting values of the cooperatively regulated and PSI-matched control events were used as input for a two-sample Wilcoxon test. Resulting *P* values were corrected for multiple hypothesis testing using the Benjamini-Hochberg procedure. Windows with an FDR ≤ 0.01 were considered as having a significant signal difference.

### Sequence composition of HNRNPH binding sites

For each of the 468,452 binding sites the nucleotide sequence in a window of ± 25-nt around the center was extracted and used to create a logo plot reflecting the nucleotide composition around HNRNPH binding sites.

Tetramer frequencies in a ± 25-nt window around the binding site centers were analyzed in two pairwise comparisons. Once between strong and weak binding sites and once between binding sites located in cooperatively regulated and control events. Binding sites were considered as strong if they belong to the 20% of binding sites with the highest PureCLIP score and weak if they belong to the 20% of binding sites with the lowest PureCLIP score. For each of the four groups of binding sites, the mean frequency of each tetramer in the 51-nt windows was calculated and used for scatter plots. In **Figure 2E**, points are colored according to the presence of a G-duplet or G-triplet in the tetramer and the 12 most abundant tetramers are highlighted.

### RNA G-quadruplex prediction

RNA G-quadruplexes (rG4s) were predicted in a ± 50-nt window around the center of all binding sites and in the region of regulated and control cassette exon events using the R/Bioconductor package pqsfinder (version 2.9.1) (Hon et al. 2017) with the following parameters: strand = “+”, max_len = 30, min_score = 25, loop_min_len = 1, loop_max_len = 7 and max_defects = 0. The region of cassette exon events was defined as the first position of the upstream exon to the last position of the downstream exon.

### Dependence of binding and cooperativity on rG4s

The 5-nt HNRNPH binding sites were split into four groups based on the presence of a G-triplet and the overlap with a predicted rG4. The four groups were compared for their PureCLIP score to check for a dependence of HNRNPH binding on G-runs and rG4 formation.

To examine the relationship in the degree of cooperativity and rG4s, we first determined the number of predicted rG4s overlapping the alternative exon and the flanking intronic regions of cooperatively enhanced, repressed and control cassette exon events. The intronic regions include the last up to 300 nt of the upstream intron as well as the first and last up to 300 nt of the downstream intron. The beginning of the upstream intron was not considered as the RNA splice maps revealed no enriched binding in this region. Fraction of events with a certain number of predicted rG4s overlapping the alternative exon and the intronic regions was compared afterwards. In addition, cooperatively enhanced and repressed events were split each into five groups by increasing Hill coefficient. Groups were afterwards compared for differences in the fraction of events with a certain number of rG4s overlapping the alternative exon (repressed events) or the intronic regions (enhanced events).

### Processing of RTstop profiling data

After running basic quality checks using FastQC (version 0.11.8) (https://www.bioinformatics.babraham.ac.uk/projects/fastqc/), RTstop profiling reads were filtered based on sequencing qualities (Phred score) in the sample barcode and UMI regions using the FASTX-Toolkit (version 0.0.14) (http://hannonlab.cshl.edu/fastx_toolkit/) and seqtk (v1.3) (https://github.com/lh3/seqtk/). All read pairs with a Phred score below 10 in the barcode or UMI regions of R1 were removed from further analysis. Read pairs were de-multiplexed based on the sample barcode, which is found on positions 6 to 11 of the first read (R1), using Flexbar (Roehr et al. 2017) (version 3.4.0). Subsequently, barcode and UMI regions as well as adapter sequences were trimmed from read ends of R1 using Flexbar requiring a minimal overlap of 1-nt of read and adapter. UMIs were added to the read identifiers of R1 and R2. Reads shorter than 15-nt were removed from further analysis. Using basic Bash commands and seqtk (version 1.3), R1 reads of each sample were de-multiplexed further into one FASTQ file per oligonucleotide per sample based on the oligonucleotide barcode, which is found in the first 15-nt of R2.

Reads (R1) of each oligonucleotide were mapped to the respective olionucleotide sequence using STAR (version 2.6.1b) (Dobin et al. 2013). When running STAR, up to 4% mismatched bases were allowed, soft-clipping was prohibited on the 5’ end of reads and only uniquely mapped reads were kept for further analysis. The position upstream of each read mapped to the forward strand was extracted and summarized to bedGraph and bigWig tracks using the BEDTools suite ((Quinlan and Hall 2010), version 2.27.1) and bedGraphToBigWig of the UCSC tool suite (version 365) (https://github.com/ENCODE-DCC/kentUtils/).

To quantify the number of rG4s in the IVTs, a custom RTstop peak-calling approach was implemented to identify “RTstop peaks”, i.e. 3-nt windows with RTstop signal above background. In brief, peak calling was performed independently for each buffer condition (KCl and NaCl), on the RTstop signal (read starts) summed across all three replicates per condition. To account for read start enrichment at the 5’ end of constructs lacking rG4 formation—i.e., when reverse transcription is not hindered by secondary structure—the first 18 nt of the 200-nt constructs were excluded from the seed position search. Subsequently, positions 19 to 200 were scanned for positions containing at least six stacked RTstop read starts, which were defined as seed sites. Seed sites were further processed using BindingSiteFinder (version 1.3.8) (Busch et al. 2020) to generate 3-nt wide RTstop peaks, requiring that at least two out of three positions were supported by RTstop signal (bsSize = 3, minWidth = 1, minCrosslinks = 2, minCLSites = 1). Peaks from both conditions were then merged, and partial overlaps of peaks were resolved by resizing them to their center point, based on the maximum combined coverage.

To identify rG4s—i.e., RTstop peaks significantly reduced in NaCl condition—differential testing on the RTstop signal level at RTstop peaks between the conditions was performed using DESeq2 (version 1.37.2) (Love et al. 2014). Specifically, a condition-specific comparison between NaCl and KCl buffer conditions was conducted (alpha = 0.05, Benjamini-Hochberg correction). Prior to differential testing, constructs were filtered out based on the following quality criteria: (1) low expression (<100 normalized reads), (2) high variability in total reads per construct between conditions (adjusted *P* value < 0.05 in a separate DESeq2 test using the parameters altHypothesis = ‘lessAbs’ and lfcThreshold = 1). The resulting log_2_-transformed fold-changes (L2FC) were shrunken using the adaptive shrinkage method (‘ashr’). Significantly changing RTstop peaks (adjusted *P* value < 0.05) with a shrunken L2FC < 0 were considered as rG4, and constructs with at least one rG4 were labelled as rG4-containing. Finally, given that each tested transcript region was represented by four oligonucleotide constructs with distinct barcodes, only regions with two or more rG4-containing constructs were classified as rG4 regions (*n* = 1,060), while a classified non-rG4 region (*n* = 1,109) was not allowed to have a single construct labelled as rG4-containing (**Supplementary Table S5**). For each classified region, the construct with the highest read count and the respective classification was chosen as representative for all following analysis. The remaining 831 transcript regions were inconsistent and removed from further analysis.

Since many constructs harbored more than one rG4, the rG4 with the strongest RTstop signal, i.e., the highest ratio of read starts within the peak over the total number of reads on the construct, was chosen as representative peak for motif analysis (**Figure S4C**). The sum of this ratio from all rG4s on a given constructs denotes the G4 propensity of the construct. This sums is then divided by the number of peaks for normalization and give the *total construct ratio*.

### in vitro RNA library *in vitro* iCLIP profiling data processing

*In vitro* iCLIP profiling data were filtered, de-multiplexed and mapped in the same way as done and described for RTstop profiling data. Likewise, upstream positions of forward strand reads were extracted and summarized to bedGraph and bigWig tracks using the BEDTools suite (version 2.27.1) and bedGraphToBigWig of the UCSC tool suite (version 365).

### Expression and splicing quantification of TCGA BRCA samples

Read counts of genes and junction counts of BRCA samples were accessed via the R/Bioconductor package recount3 (version 1.3.9) (Wilks et al. 2021), using either “gene” or “jxn” as type in the create_rse() function, which generated a RangedSummarizedExperiment object. The samples were filtered for type “Primary Tumor” (referred to as tumor) or “Solid Tissue Normal” (referred to as normal) and “female” as participant gender. In addition, participants with more than one sample of the same type were removed (e.g., two samples labeled as tumor). The filtering resulted in 1,177 samples, including 1,069 from tumor and 108 from normal tissue.

Read counts of genes were created using compute_read_counts(), followed by normalization and transformation using the vst() function of the R/Bioconductor package DESeq2 (version 1.33.4)(Love et al. 2014). To account for variation due to processing in different batches, the normalized and transformed counts were batch-corrected using removeBatchEffect() of the R/Bioconductor package limma (version 3.49.4) (Ritchie et al. 2015) resulting in the final transcript level estimates used for correlation and clustering analyses.

Junctions stored in the RangedSummarizedExperiment object were used to calculate the PSI of regulated and control cassette exon events as well as regulated alternative last exon events. For this purpose, two junctions were defined for each of the events, namely the representative junction from the MAJIQ analysis step and the competing junction from the same LSV. In the case of cassette exon events, the competing junction is C1_C2 if C1_A is representative or C2_C1 if C2_A is representative, while for alternative last exon events the junction is called Distal. An important step before calculating the PSI values was to offset the start and end positions by 1-nt to the right and left, respectively, as the recount3 package anchors junctions at the start and end of the intron. For each event in each sample, the counts of the two junctions were extracted and, if the sum of counts was ≥ 25, used to calculate a PSI for the inclusion events by dividing the counts of the inclusion junction (C1_A or C2_A for cassette exon events and Proximal for alternative last exon events) by the sum of the counts of both junctions. Events quantified in less than 10 of the 1,177 samples were immediately removed.

In addition to the mentioned regulated and control events, PSI values of an *HNRNPH1* cassette exon (located on the minus strand of chr5), whose skipping leads to an NMD isoform, were calculated. For the calculation, the counts of the two inclusion junctions (chr5:179,619,407-179,620,892:- and chr5:179,618,323-179,619,269:-) and the skipping junction (chr5:179,618,323-179,620,892:-) were extracted for all samples and PSI values were computed by dividing the sum of the two inclusion junction counts by that value plus two times the counts of the skipping junction.

### Correlation and cluster analyses of TCGA BRCA samples

*HNRNPH1* and *HNRNPH2* transcript levels were correlated with the PSI values of cooperatively enhanced and repressed cassette and alternative last exon events and control cassette exon events, followed by multiple hypothesis testing correction of the *P* values using the Benjamini-Hochberg procedure. Correlation coefficients of the cooperatively enhanced, repressed and control events were afterwards compared.

Before clustering the 1,177 samples using the PSI values of cassette and alternative last exon events, all events quantified in less than 100 samples or having less than 10 different PSI values across all samples (e.g., all samples have full inclusion) were removed, leaving 302 events. As some samples lack quantification for an event, missing values were imputed using the imputePCA() function of the R package missMDA (version 1.18) (Josse and Husson 2016) followed by a row-wise z-score transformation. The rows and columns of the matrix were clustered using *k*-means clustering (*k* = 5). The heatmap was prepared using the R/Bioconductor package ComplexHeatmap (version 2.9.4) (Gu, Eils, and Schlesner 2016) and contains the cluster information from the *k*-means clustering as well as additional row and column annotations. One of these annotations is the subtype (Normal, LumA, LumB, Her2 and Basal), which relies on the PAM50 signature and was accessed via the PanCancerAtlas_subtypes() function of the R/Bioconductor package TCGAbiolinks (version 2.22.4) (Colaprico et al. 2016).

The capability of the cooperatively regulated events to distinguish normal from tumor samples and the Basal subtype samples from the other samples was quantified using the adjusted Rand Index calculated with the adjustedRandIndex() function of the R package mclust (version 5.4.10) (Scrucca et al. 2016). Briefly, for the distinction of normal from tumor samples, the cluster with the highest number of normal samples was determine and labeled as “C_Normal”, while the other clusters were combined into “C_Other”. The adjusted Rand Index was computed taking the new cluster labels (C_Normal and C_Other) and the sample type (normal and tumor) as class labels. To distinguish the Basal subtype samples from the other samples, the cluster with the highest number of Basal subtype samples was determined and labeled as “C_Basal”, while the other clusters were combined into “C_Other”. In addition, the non-Basal subtypes were combined as “Other”. The adjusted Rand Index was computed taking the new cluster labels (C_Basal and C_Other) and the subtype (Basal and Other) as input. A Rand Index of 0 indicates a random partition, while a value 1 indicates a perfect agreement.

To test whether splicing in the samples is generally impaired and therefore any random set of splicing events would result in a clear distinction between normal and Basal subtype samples, we picked 100 sets of control cassette exon events with a similar set size (*n* = 302) and performed the same steps as described above, i.e., imputation of missing values, *k*-means clustering (*k* = 5) and calculation of the two adjusted Rand Indices. The two adjusted Rand Indices were used to compare the 100 random sets of control events against the set of cooperatively regulated events.

### Kinetic modeling of splicing decision making

In the model depicted Fig. 6A, we assumed that splicing regulation by HNRNPH occurs in a cascade of consecutive biochemical events, i.e., cooperative HNRNPH binding to pre-mRNA (step 1), repression of multi-step spliceosome assembly by bound HNRNPH (step 2), splicing decision making based on spliceosome binding (step 3). As described in the **Supplemental Material**, the following equations describe the three cascade levels (i)

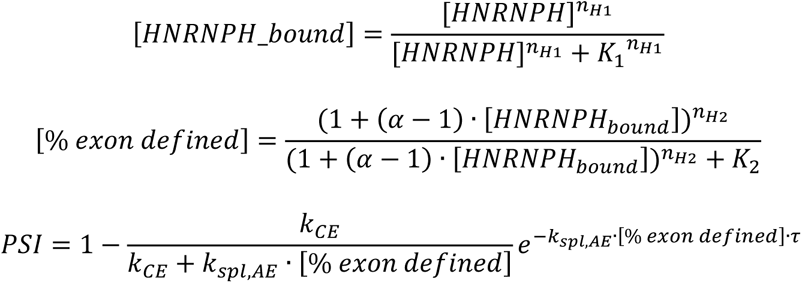

To simulate dose-response curves, the HNRNPH concentration (input) was over a broad concentration range with logarithmic spacing. The Hill coefficients of the individual levels and the overall Hill coefficient of the cascade was quantified using the formula (ii)

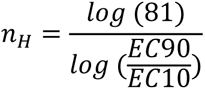

Here, EC10 and EC90 are the input concentrations that lead to 10% and 90% of the maximal output. For instance, for the overall cascade dose-response, the input was the HNRNPH concentration, and the output was the PSI. Before applying Eq. ii, each dose-response curve was min-max-normalized between 0 and 1. In **Figure 6D and Figure S7C**, 1000 simulation runs were performed, in which each of the parameters was independently sampled from the distributions indicated (see **Supplemental Material**).

### Analysis of rG4-disrupting genetic variants in breast cancer patients

Genetic single nucleotide variants (SNVs) were obtained from the Penn Medicine BioBank (PMBB), including breast cancer patients of European (*n* = 29,362) and African (*n* = 10,217) ancestry. Variants were filtered to exclude singletons and those with high missing call rates (>10%). Samples were excluded based on low exome sequencing coverage (below 85% of targets reaching 20x coverage), high contamination (D-stat > 0.4), genetically identified sample duplicates, and sex mismatches. Only second-degree or more unrelated individuals were included within each ancestry group. A total of 2,664,044 filtered SNPs were considered in the following analyses. All genomic coordinates and variant predictions were based on the GRCh38/hg38 human genome assembly.

Using G4mer (https://huggingface.co/Biociphers/g4mer), (Zhuang et al. 2024) we predicted RNA G-quadruplexes (rG4s) in all genomic 70-nt windows overlapping HNRNPH binding sites, considering the maximum G4mer score for each sequence. This yielded 173,950 binding sites containing at least one predicted rG4 (G4mer score > 0.5). Comparison with PMBB data identified 51,562 SNVs that overlapped with the predicted rG4 windows at HNRNPH binding sites. For each SNV, we substituted the alternative allele into the 70-nt predicted rG4 sequence and rescored it using G4mer. Variants were classified as rG4-disrupting if (i) the G4mer score dropped by more than 0.2 (Δ score < –0.2) and (ii) the alternative sequence score was below 0.5. In total, we identified 13,482 rG4-disrupting PMBB variants in the vicinity of HNRNPH binding.

We then tested 13,473 of these variants for association with breast cancer diagnosis, using logistic regression models adjusted for sex, age at enrollment, age-squared, and the first 10 principal components (PCs) of genetic ancestry. Breast cancer diagnoses were defined based on ICD-10 codes mapped to phecodes using the PheWAS package (v 0.99.6.1, (Carroll, Bastarache, and Denny 2014)) in R, and patients were considered ‘cases’ if they had at least two occurrences of relevant codes on different days, improving phenotype precision. This analysis yielded 8 rG4-disrupting variants significantly associated with breast cancer (false discovery rate, q < 0.05). Of these, 6 SNVs were observed only in patients of European ancestry, and 2 SNVs were observed only in African ancestry.

## Data availability

The sequencing data generated in this study have been deposited in the NCBI Gene Expression Omnibus (GEO) database under Series accession GSE303138 (https://www.ncbi.nlm.nih.gov/geo/query/acc.cgi?acc=GSE303138), including RNA-seq (GSE303137), iCLIP (GSE303135), RTstop profiling (GSE304967) and in vitro iCLIP (GSE304723). The secure token for anonymous reviewer access is: **mdatwgsktrwxpqz**

## Code availability

The computational code for the analyses and figure generation is available at https://github.com/ZarnackGroup/Publications/tree/main/Tretow_et_al_2025.

## Acknowledgements

We are grateful to all members of the participating labs for their support and discussion. We thank Jean-Louis Mergny for initial experiments and discussion. Support by the IMB Protein Production Core Facility for the purification of recombinant HNRNPH1 is gratefully acknowledged. We gratefully acknowledge the Institute of Molecular Biology (IMB) Core Facilities for their support, especially the Genomics Core Facility and the use of its NextSeq 500 (funded by the Deutsche Forschungsgemeinschaft [DFG, German Research Foundation] P#329045328) and the Bioinformatics Core Facility. The results published here are in part based upon data generated by the TCGA Research Network: https://www.cancer.gov/tcga.

## Author contributions

K.T. performed most experiments, including transcriptomics, *in vitro* binding studies and reporter assays. M.K. and M.B. performed bioinformatics analyses with help of J.B. and A.B.. M.M., M.C., N.K., S.B., N.H., N.M., and H.H. supported experiments. S.L. and F.S. performed modeling. F.Z. and Y.B. performed genetic variant analyses. P.B. and S.S. contributed ideas and reagents. K.T., M.K., K.Z., and J.K. designed the study. K.Z. and J.K. supervised bioinformatics analyses and experimental work, respectively. K.T., M.K., M.C., K. Z., and J. K. wrote the manuscript with help and comments from all co-authors.

