## Supplemental File for "RNA G-quadruplexes mediate cooperativity in HNRNPH binding and splicing regulation"

### Supplementary Methods

#### Model of multi-step splicing regulation by HNRNPH

In the model depicted in **Figure 6A**, we assumed that splicing regulation by HNRNPH occurs in a cascade of consecutive biochemical events (steps 1–3). Here, we derive equations describing each level, analytically calculate accessible Hill coefficients and provide parameter values used in the simulations in **Figure 6D** and **Figure S7C**.

***Step 1 – Cooperative HNRNPH binding to pre-mRNA***

The first step in the splicing-regulatory cascade is HNRNPH binding to pre-mRNA. In line with the *in vitro* titration experiments (**Figure 4C–J**), we assume HNRNPH binding to be cooperative, and describe it using a Hill equation with a Hill coefficient of $n_{H1}$ (1)

$$[HNRNPH\_bound] = \frac{[HNRNPH]^{n_{H1}}}{[HNRNPH]^{n_{H1}} + {K_1}^{n_{H1}}}$$

For high HNRNPH concentrations, the binding equation reaches a value of 1, reflecting full occupancy of all pre-mRNA molecules with HNRNPH. The parameter $K_1$ describes the HNRNPH concentration at which the binding is half-maximal and reflects the affinity of HNRNPH for pre-mRNA.

***Step 2 – HNRNPH effect on spliceosome binding***

The second step in the splicing-regulatory cascade is the binding of pioneering spliceosomal subunits (U1, U2) to splice sites of the alternative exon. In our model, we assume an exon definition mechanism, in which U1 and U2 cooperatively bind across an exon (J. E. Braun et al. 2018; De Conti, Baralle, and Buratti 2013) as described in the literature and shown in **Figure S7A**, we assume that U1 is bound first to the pre-mRNA and that the resulting pre-U1 complex recruits U2. If we assume U1 and U2 to be present in excess over pre-mRNA the differential equations of this model are given by (2)

$$\frac{d[premRNA]}{dt} = -k_{on,U1} \cdot [premRNA] \cdot [U1] + k_{off,U1} \cdot [U1 - pre]$$

$$\frac{d[U1U2 - pre]}{dt} = k_{on,U2} \cdot [U1 - pre] \cdot [U2] - k_{off,U2} \cdot [U1U2 - pre]$$

With the mass conservation relation (3)

$$[premRNA] + [U1 - pre] + [U1U2 - pre] = 1$$

Setting the differential equations to zero, we derive the following equilibrium binding equation (4)

$$[\%\, exon\, defined] = [U1U2 - pre] = \frac{k_{on,U1} \cdot k_{on,U2} \cdot [U1] \cdot [U2]}{k_{on,U1} \cdot [U1] \cdot \left(k_{on,U2} \cdot [U2] + k_{off,U2}\right) + k_{off,U1} \cdot k_{off,U2}}$$

For high cooperativity, i.e., very strong U2 binding to the U1-pre-spliceosomal complex ($k_{on,U2} \cdot [U2] \gg k_{off,U2}$), this simplifies to (5)

$$[\%\, exon\, defined] = [U1U2 - pre] = \frac{k_{on,U1} \cdot k_{on,U2} \cdot [U1] \cdot [U2]}{k_{on,U1} \cdot k_{on,U2} \cdot [U1] \cdot [U2] + k_{off,U1} \cdot k_{off,U2}}$$

HNRNPH is assumed to regulate splicing by lowering the affinity U1 and/or U2 binding, acting as a repressor of alternative exon definition. If a bound HNRNPH molecule simultaneously affects both U1 and U2 binding, we can write the parameters $k_{on,U1}$ and $k_{on,U2}$ as (6)

$$k_{on,U1} = k_{on,U1}^{free} \cdot (1 - [HNRNPH_{bound}]) + k_{on,U1}^{bound} \cdot [HNRNPH\_bound]$$

$$k_{on,U2} = k_{on,U2}^{free} \cdot (1 - [HNRNPH_{bound}]) + k_{on,U2}^{bound} \cdot [HNRNPH\_bound]$$

Here, $k_{on}^{free}$ and $k_{on}^{bound}$are the on-rates in the absence and presence of HNRNPH, respectively. Rearranging yields (7)

$$k_{on,U1} = k_{on,U1}^{free} + \left(k_{on,U1}^{bound} - k_{on,U1}^{free}\right) \cdot [hnRNPh\_bound]$$

$$k_{on,U2} = k_{on,U2}^{free} + \left(k_{on,U2}^{bound} - k_{on,U2}^{free}\right) \cdot [hnRNPh\_bound]$$

When assuming that HNRNPH binding reduces the on-rate by a factor α, i.e., (8)

$$k_{on,U1}^{bound} = \alpha_1 \cdot k_{on,U1}^{free}$$

$$k_{on,U2}^{bound} = \alpha_2 \cdot k_{on,U2}^{free}$$

we obtain (9)

$$k_{on,U1} = k_{on,U1}^{free} \cdot (1 + (\alpha_1 - 1) \cdot [hnRNPh_{bound}])$$

$$k_{on,U2} = k_{on,U2}^{free} \cdot (1 + (\alpha_2 - 1) \cdot [hnRNPh_{bound}])$$

For simplicity, we presume that HNRNPH binding reduces U1 and U2 binding to the same extent ($\alpha = \alpha_1 = \alpha_2$) and using Eq. 5, we describe exon definition as a function HNRNPH binding using the following formula (10)

$$[\%\, exon\, defined] = \frac{(1 + (\alpha - 1) \cdot [hnRNPh_{bound}])^2}{(1 + (\alpha - 1) \cdot [hnRNPh_{bound}])^2 + K_2}$$

where $K_2$ is a lumped parameter (11)

$$K_2 = \frac{k_{off,U1} \cdot k_{off,U2}}{k_{on,U1}^{free} \cdot k_{on,U2}^{free} \cdot [U1] \cdot [U2]}$$

More generally, when HNRNPH affects more or less than two steps in spliceosome assembly, Eq. 10 modifies to (12)

$$[\%\ exon\ defined] = \frac{(1+(\alpha-1)\cdot[HNRNPH_{bound}])^{n_{H2}}}{(1+(\alpha-1)\cdot[HNRNPH_{bound}])^{n_{H2}}+K_2}$$

***Step 3 - From spliceosome binding to splicing outcomes***

After the exon definition step, the cross-exon U1-U2 complex is converted into a cross-intron U1-U2 complex to which the catalytic spliceosome subunits U4-U6 are recruited to catalyze the excision of bridged introns (De Conti et al. 2013). Since, cross-intron complexes can only form between two fully defined exons, the exon definition state of the alternative exon (AE), i.e., step 2 above, determines splicing outcomes: If the AE is defined it can form catalytic complexes with both neighboring exons (which we assume here to be constitutive), implying that the AE will be included in the final transcript (inclusion isoform). If the AE remains undefined the cross-intron complex forms between the outer constitutive exons and the AE is excluded (skipping isoform).

Baeza-Centurion et al. derived an equation describing co-transcriptional splicing decision making due to the competition of two exons (Baeza-Centurion et al. 2019): (i) an upstream alternative exon (exon A in **Figure S7B**) and a downstream constitutive exon (exon B in **Figure S7B**). They described the percent splice-in (PSI) metric as a function of the usage rate of the two exons, reflected by the two parameters, $k_{AE}$ and $k_{CE}$ (13)

$$PSI = 1 - \underbrace{\frac{k_{CE}}{k_{CE}+k_{AE}}}_{term\ I}\underbrace{e^{-k_{AE}\cdot\tau}}_{term\ II}$$

According to term I, which represents purely post-transcriptional splicing, the two splicing outcomes compete, i.e., skipping will be the primary outcome (PSI≈0) if the CE usage rate is much higher than the AE usage rate ($k_{CE} >> k_{AE}$), whereas high inclusion is observed in the opposite scenario. Term II contains a time delay τ that reflects the co-transcriptional nature of splicing decisions: for slow transcript elongation (high τ) inclusion is the preferred splicing outcome irrespective of the exact choice of $k_{CE}$ and $k_{AE}$ as commitment to inclusion it is possible earlier during transcript elongation (see scheme in **Figure S7B**).

In our model, we assume the exon usage parameter of the AE to be proportional to the definition state of that exon (14)

$$k_{AE} = k_{spl,AE}\cdot[\%\ exon\ defined]$$

where the parameter $k_{spl,AE}$ reflects the speed of formation of splicing-committed spliceosome complexes towards inclusion after the exon has been defined.

***Hill coefficient of the full splicing-regulatory cascade***

To describe how changes in the cellular HNRNPH concentration affect splicing outcomes, we combine Eqs. 1, 12 and 13, i.e., the full model reads (15)

$$[hnRNPh\_bound] = \frac{[hnRNPh]^{n_{H1}}}{[hnRNPh]^{n_{H1}} + K_1{}^{n_{H1}}}$$

$$[\%\, exon\, defined] = \frac{(1 + (\alpha - 1) \cdot [hnRNPh_{bound}])^{n_{H2}}}{(1 + (\alpha - 1) \cdot [hnRNPh_{bound}])^{n_{H2}} + K_2}$$

$$PSI = 1 - \frac{k_{CE}}{k_{CE} + k_{spl,AE} \cdot [\%\, exon\, defined]} e^{-k_{spl,AE} \cdot [\%\, exon\, defined] \cdot \tau}$$

***Choice of parameter values:*** Eq. 14 was used to simulate the splicing-regulatory cascade in **Figure 6B–D**. The following table lists the choice of all parameter values in each simulation.

| | **Figure 6B–C** | **Figure 6D and S7C** |
|---|---|---|
| $K_1$ | 1 | Log-normal distribution (μ=1, σ=0.7) |
| $n_{H1}$ | 2 | Normal distribution (μ=2, σ=0.4) |
| $K_2$ | 0.5 | Log-normal distribution (μ=1, σ=0.7) |
| $n_{H2}$ | 2 | Normal distribution (μ=2, σ=0.4) |
| α | 0.01 | Log-normal distribution (μ=1, σ=0.7) |
| $k_{CE}$ | 0.1 | Log-normal distribution (μ=0.02, σ=0.7) |
| $K_{spl,AE}$ | 2 | Log-normal distribution (μ=0.02, σ=0.7) |

In all simulations, the HNRNPH concentration (input) was over a broad concentration range with logarithmic spacing. The Hill coefficient of the PSI was quantified using the previously established formula (Huang and Ferrell 1996)(16)

$$n_H = \frac{log\,(81)}{log\,(\frac{EC90}{EC10})}$$

Here, $EC_{10}$ and $EC_{90}$ are the input (HNRNPH) concentrations that lead to 10% and 90% of the maximal output (PSI). If the output curve minima and maxima deviating from 0 and 1, respectively, it was rescaled between 0 and 1 before applying Eq. 16.

In **Figure 6D** and **S7C**, 1,000 simulation runs were performed, in which each of the parameters was independently sampled from the distributions indicated above.

### Quantitative biophysical modelling

We developed a quantitative biophysical model to describe HNRNPH binding events to folded and unfolded rG4 structures. We describe the system by three characteristic free energies: (i) the free energy difference between a folded rG4 structure and an unfolded one, $g_{\mathrm{rG4}}$, (ii) the binding free energy for a single binding event, $g_{\mathrm{bind}}$, (iii) an entropic loop penalty, $g_{\mathrm{backfold}}$, if an HNRNPH protein loops back to bind once more. To keep the model simple, we do not distinguish between different types of loops and RRM domains. We use the following notation: $0^F$ for the folded rG4 state, $0$ for the unfolded state with no bound proteins, $j^{b_1 b_2 \ldots b_j}$ for binding states with $j$ bound proteins, where $b_i$ is the number of bonds between the rG4 sequence and the protein $i$. Selected examples of binding states are shown in (**Figure 4L**). Taking the state $0$ as reference state, the relative concentrations of rG4 structures in the binding state $j^{b_1 b_2 \ldots b_j}$ at equilibrium is

$$\frac{c_{j^{b_1 b_2 \ldots b_j}}}{c_0} = \rho_p^j \, \frac{m!}{j!\,(m-b!)} \left(n\, e^{g_{\mathrm{bind}}/RT}\right)^j \prod_{i=1}^{j} \left[ e^{(b_j-1)(g_{\mathrm{bind}}-g_{\mathrm{backfold}})/RT} \left(n-1\; b_j-1\right) \frac{1}{b_j} \right],$$

where $m = 4$ is the number of available binding sites on the rG4 structure, $n = 3$ is the number of available RRM domains, $b = \sum b_j$ is the total number of occupied binding sites on the RNA in the state $j^{b_1 b_2 \ldots b_j}$, $\rho_p$ is the protein concentration in solution, and we have accounted for the different possibilities to distribute $(b_1, .., b_j)$ bonds between proteins and RNA binding sites by a combinatorial factor. Furthermore, we have $\frac{c_{0^F}}{c_0} = e^{g_{\mathrm{rG4}}/RT}$

From these expressions, we can derive the fraction $f(\rho_p) = c_{\mathrm{bound}}/c_{\mathrm{total}}$ of rG4 structures which are bound to at least one protein with respect to the total concentration of rG4 structures. This is then used to derive the transition density or activity coefficient $K$ via $f(K) \coloneqq 1/2$ and the Hill coefficient $n$ via $n_H = 2Kf'(K)/f(K)$.

To analyze the behavior of the model, we first considered two extreme cases: One where only binding states without backfolding are allowed (shown inside the dashed box in **Figure 4K**), and one where loops are not penalized ($g_{\mathrm{backfold}} = 0$). The first case typically results in a Hill coefficient of $n_H = 4$ for sufficiently large binding and folding energies (**Figure 4L**), whereas the second case results in $n_H = 2$ for a broad parameter range (**Figure 4M**). Importantly, one generally has $n_H \leq 2$ if the activation coefficient is below $K < 0.1$ mol/l, independent of the binding energy, $g_{\mathrm{bind}}$. A closer inspection revealed that the behavior of systems with $n_H \approx 2$ is dominated by two binding states, $2^{31}$ and $2^{22}$, highlighted in (**Figure 4K**), where two proteins bind with two loops such that all four RNA sites are occupied.

Next we used the experimental data to further refine the model. The data suggest an activity coefficient $K \approx 3 \cdot 10^{-6}$mol/l, which constrains the combinations of possible free energy parameters, $g_{\mathrm{rG4}}$, $g_{\mathrm{bind}}$, and $g_{\mathrm{backfold}}$. For instance, if one neglects all binding states except the dominant two, $2^{31}$ and $2^{22}$("two-loop binding only" model), one gets the simple relation $RT\ \ \ln(12\,K)\ = \frac{1}{2}g_{\mathrm{rG4}} - 2g_{\mathrm{bind}} + g_{\mathrm{backfold}}$. For $K = 3 \cdot 10^{-6}$mol/l, this equation also describes the data for the full model reasonably well for all $g_{\mathrm{backfold}} < 7$ kcal/mol (**Figure S5E**). For higher $g_{\mathrm{backfold}}$, deviations set in, indicating that the loop penalty increasingly suppresses the two dominant binding states, $2^{31}$ and $2^{22}$, as also indicated by an increase of the Hill coefficient from $n_H = 2$ to $n_H = 4$ (**Figure S5F**). Since we have measured $n_H \approx 2$ (ARPC2), we concluded that $g_{\mathrm{backfold}}$ must be small. The folding energy of rG4 was measured by Yu et al (Yu et al. 2009) as $g_{\mathrm{rG4}} \approx 23$ kcal/mol. The binding energy for RNA/protein interactions is estimated as $g_{\mathrm{bind}} = 9.2$ kcal/mol from the molecular dissociation constant $K_d = 0.2\ \cdot 10^{-6}$ mol/l of RRM domains of closely related proteins and G-tract RNAs as measured by Tamayo et al (Tamayo et al. 2017). Inserting this value, we conclude that the loop penalty, $g_{\mathrm{backfold}}$ must be of order 0.8 kcal/mol.

Finally, we also investigated the impact of admitting a helper state, $1^F$, where a single RRM domain binds to folded rG4 with binding energy $g_{\mathrm{rG4,bind}}$, in order to assist unfolding. As long as $g_{\mathrm{rG4,bind}}$ is smaller than 4 kcal/mol, we found that the presence of such a state has no significant influence on the results. However, if the binding energies to the folded and unfolded rG4 structures are comparable, then the quadruplex binding dominates and the cooperative behavior disappears, i.e., $n_H = 1$ (**Figure 4O**).

### Supplementary Figures

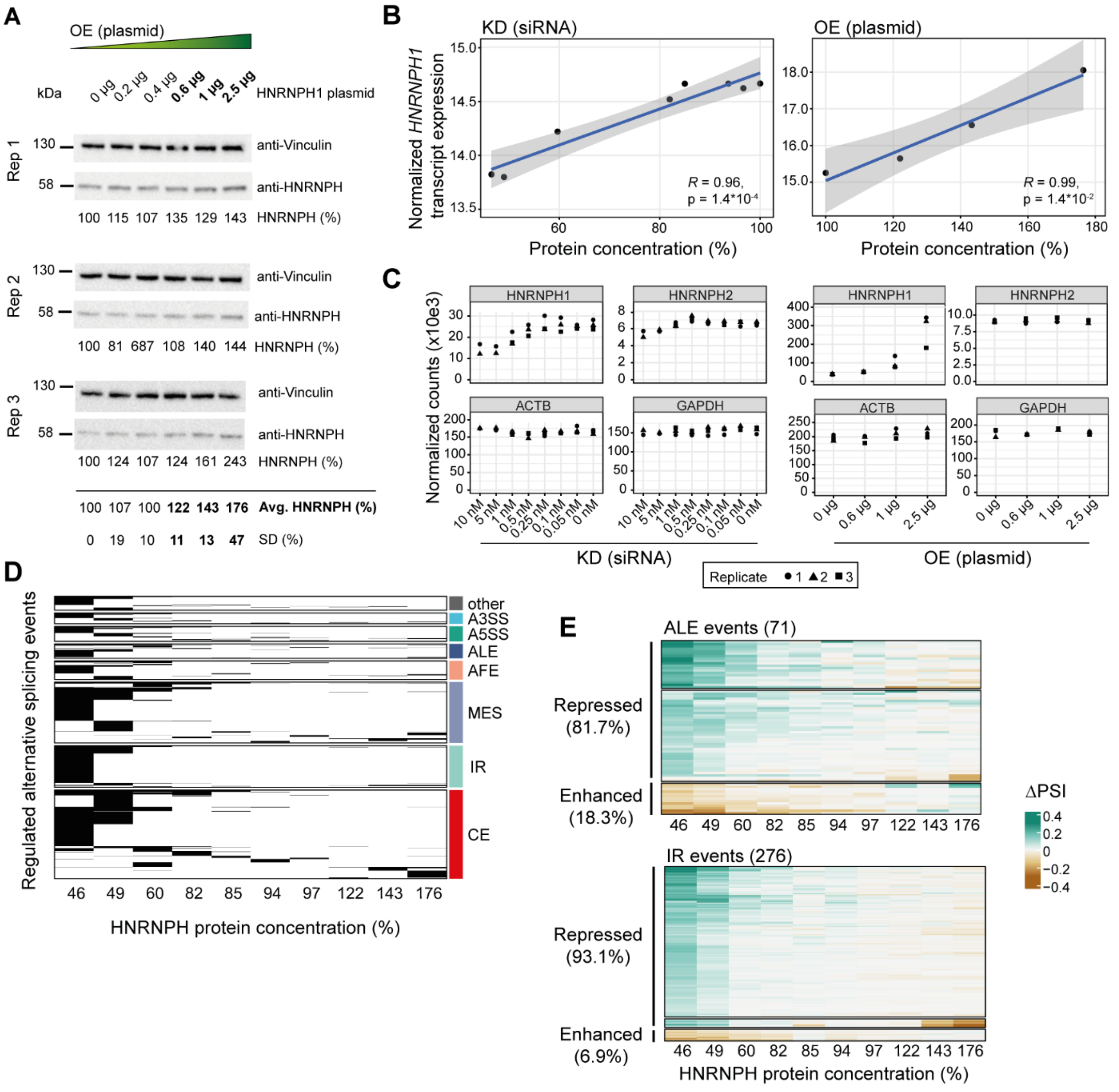

**Supplement Figure S1. Titration of HNRNPH leads to changes in alternative splicing. (A)** Western blot for HNRNPH and Vinculin, measuring overexpression on the protein level for three biological replicates. Mean and standard deviation are indicated on the plot. **(B)** Scatter plots show the correlation between relative HNRNPH protein concentration and normalized *HNRNPH1* transcript expression (vst-transformed RNA-seq read counts, comparable to $\log_2$ scale) in knockdown (left) and overexpression (right) experiments. Pearson correlation coefficients (*R*) and the associated *P* values are indicated. **(C)** Normalized RNA-seq read counts of *HNRNPH1*, *HNRNPH2*, *ACTB* and *GADPH* across a series of *HNRNPH1* KD (siRNA; 10–0 nM) and overexpression (plasmid; 0–2.5 µg) conditions. **(D)** Heatmap visualization shows the regulation of alternative splicing events across a range of HNRNPH protein concentrations (46–176%). Conditions in which an alternative splicing event was significantly changed are shown in black. **(E)** Heatmap shows changes in inclusion levels (ΔPSI) of ALE (rows; upper plot), and IR (rows; lower plot) across range of HNRNPH protein concentration (46–176%). Events shown were quantified in all 10 comparisons and regulated in at least one of the two strongest knockdown and/or overexpression conditions.

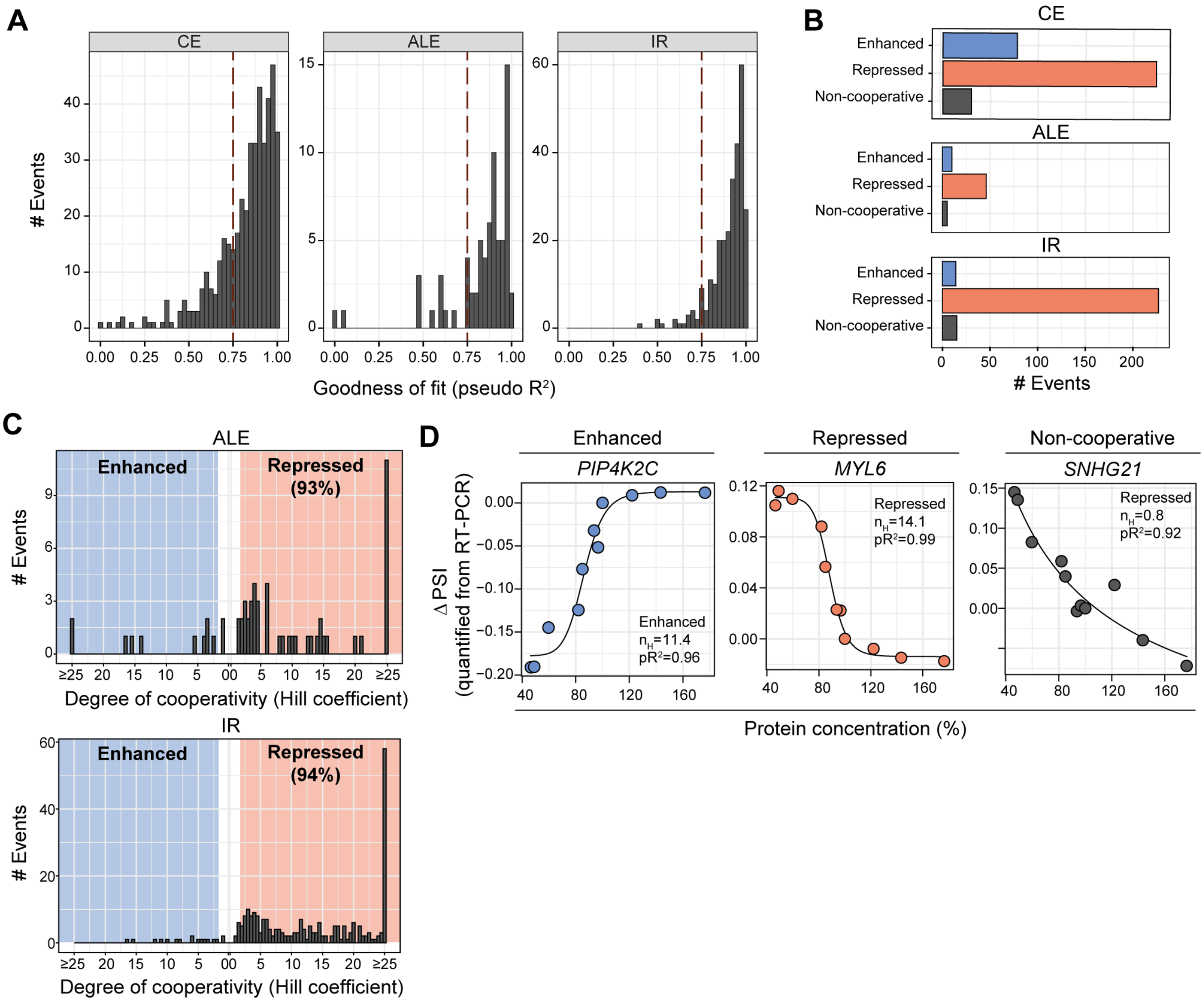

**Supplement Figure S2. Switch-like splicing response of cassette exons during titration hints towards cooperative regulation by HNRNPH. (A)** Bar plots showing the goodness of fit (pseudo $R^2$) for fitted first-order Hill equations extracted from dose-response curves for all regulated CE (left), ALE (middle), and IR (right) events. **(B)** Bar chart shows the number of CE (top), ALE (middle), and IR (bottom) events that are cooperatively regulated ($n_H \geq 2$) and enhanced or repressed, as well as events that lack cooperative regulation (Non-cooperative; $n_H < 2$). **(C)** Histograms show the distribution of Hill coefficient estimates for all HNRNPH-dependent ALE (top) and IR (bottom). Value ranges considered as cooperatively enhanced (blue) or repressed (red) are indicated. **(D)** Scatter plots show inclusion levels (ΔPSI; quantified from RT-PCR) against HNRNPH protein levels for selected examples, showing cooperatively enhanced (exon 5 of *PIP4K2C*; left), repressed (exon 6 of *MYL6*; middle) and non-cooperative (exon 3 of *SNHG21*; right) regulation. To test for cooperativity, a four-parametric logistic curve (also known as the Hill equation) was fitted to the data points (solid line). Hill coefficient ($n_H$) and pseudo $R^2$ (p$R^2$) are indicated on the plot.

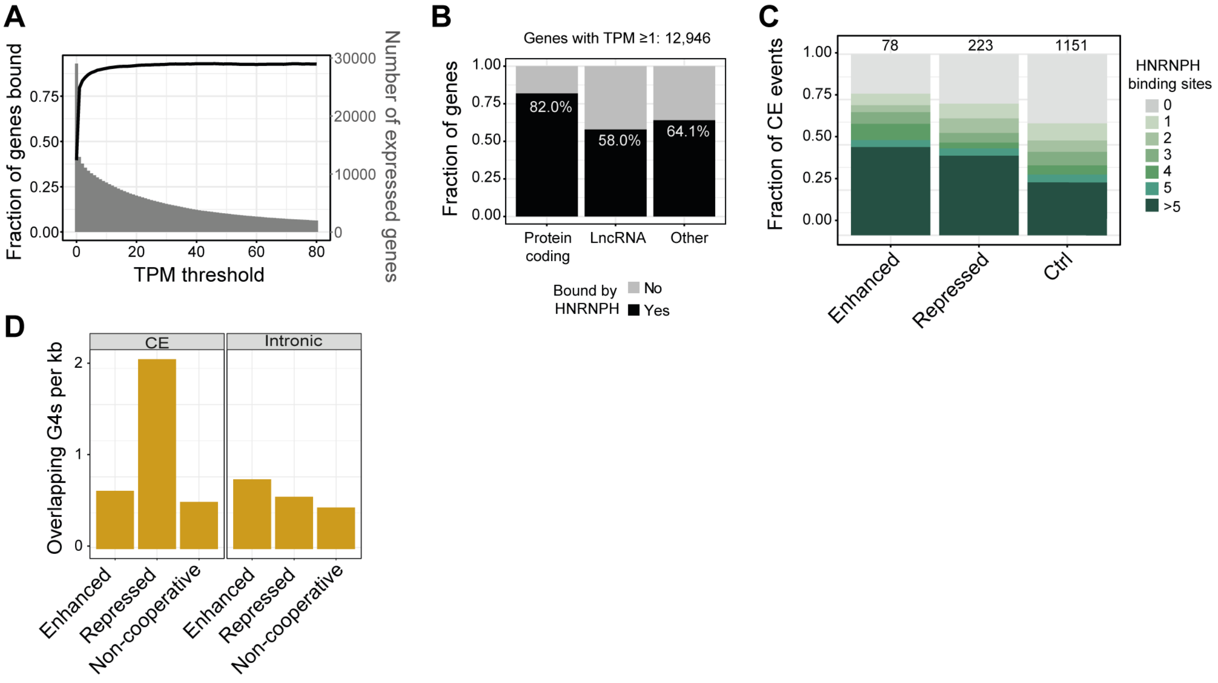

**Supplement Figure S3. The positioning of HNRNPH determines the splicing outcome. (A)** Plot shows fraction of genes bound (black) and number of expressed genes (gray) against TPM threshold (0–80). **(B)** Fraction of transcripts bound by HNRNPH (black) within protein coding genes, lncRNA and others. Number of genes included in analysis (TPM ≥ 1) is indicated. **(C)** Stacked bar plot shows distribution of cooperatively enhanced (left), repressed (middle), and control (right) CE split by number of HNRNPH binding sites they harbor (gray to green shading). **(D)** Bar plots show number of overlapping G4s for CE (left) and intronic sequences (right) that are cooperatively enhanced, repressed or control sequences.

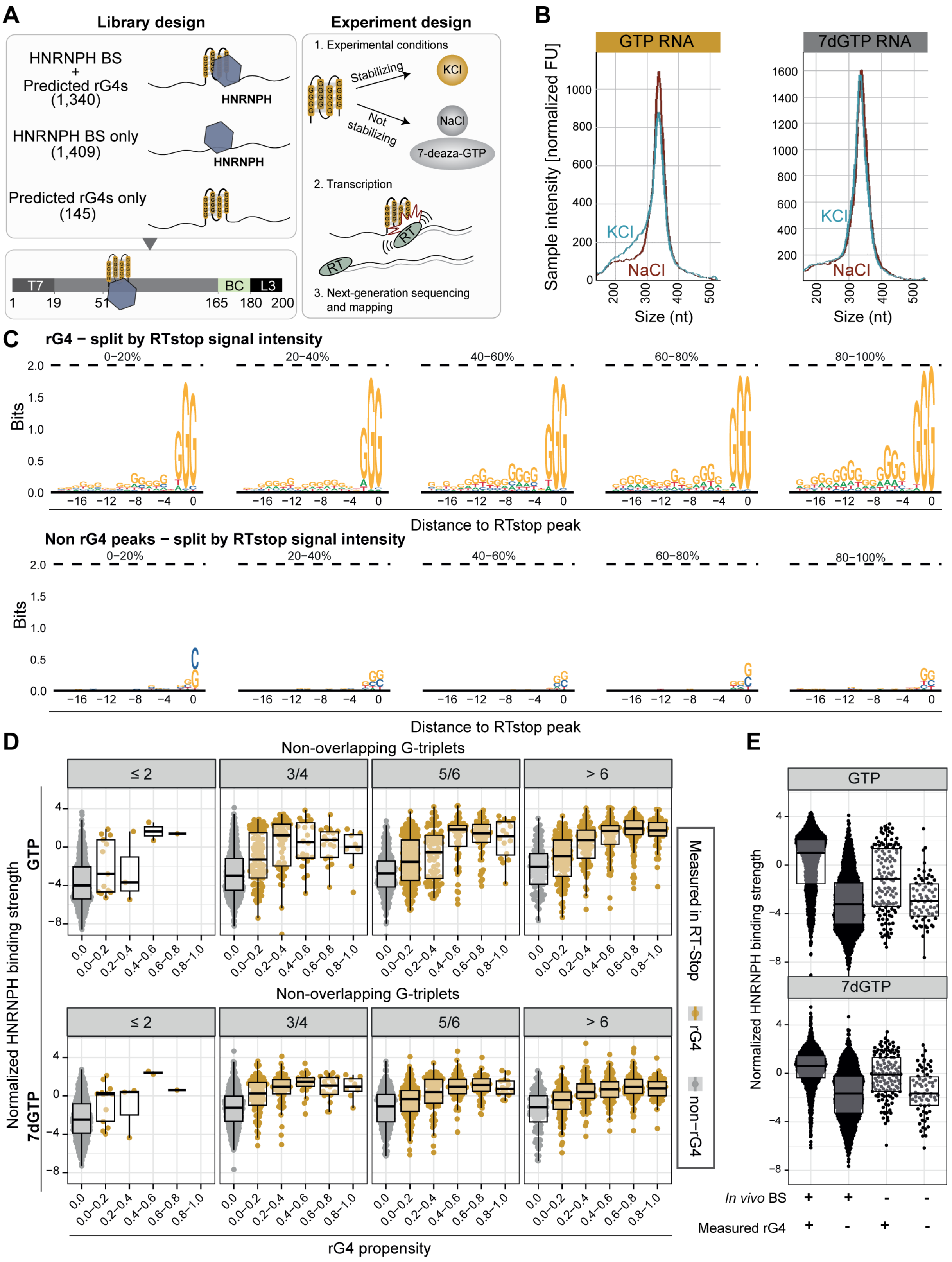

**Supplement Figure S4. HNRNPH binding sites fold into RNA rG4s *in vitro*. (A)** Schematic overview of experimental design of reverse transcriptase stop (RTstop) profiling (Guo and Bartel 2016). The protocol was adopted to using a pool of 11,576 *in vitro* transcripts (IVTs), representing 2,894 different transcript regions: 1,409 HNRNPH binding sites (BS-only), 145 predicted rG4s (rG4-only), and 1,340 predicted rG4s with an HNRNPH binding site (BS+rG4)

transcribed from a designed oligonucleotide library. Each transcript region was represented by four oligonucleotides with distinct barcode sequences. Each 200-nt DNA oligonucleotide contained 146 nt of transcript sequence, flanked by a T7 promoter for *in vitro* transcription, and an L3 linker sequence for subsequent library preparation. RT is impeded by rG4s and other strong secondary RNA structures, resulting in truncated cDNA that terminates at the RNA folds. To distinguish rG4s from other RNA folds, the RT reaction was performed in the presence of ions that do or do not stabilize rG4s (KCl or NaCl, respectively). In addition, IVTs were transcribed in presence of GTP analog 7dGTP, which cannot engage in rG4s, but does not interfere with HNRNPH binding. **(B)** Plots show sample intensity distribution for RNA synthesized with GTP (left) or 7dGTP (right) in NaCl (red) and KCl (blue) buffers. Visual inspection of the RT-stop profiling results in a genome browser confirmed that reverse transcription (RT) truncates at predicted rG4 sites in KCl but not in NaCl. However, when we used 7-deaza-GTP RNA for RTstop profiling, RT truncation in KCl was no longer detected. **(C)** Sequence logos for around rG4s (top), and non-rG4 peaks (bottom), split into 20%-percentile based on their RTstop signal intensity (read starts within peak over total reads per construct). Only the strongest rG4 in each construct was included in the analysis. **(D)** Boxplots show the normalized HNRNPH binding strength across different rG4 propensities, under conditions where IVTs are transcribed with GTP (up) and 7dGTP (down). The data is divided by non-overlapping G-triplet counts (≤2, 3/4, 5/6, >6). rG4 propensity, as predicted by RTstop, is plotted in increasing bins on the x-axis. The violin plots represent the distribution of normalized HNRNPH binding strength for non-rG4 (gray) and rG4 (yellow) elements. Box plots inside violins indicate median and interquartile ranges. **(E)** Comparison of normalized HNRNPH binding strength between GTP (upper) and 7dGTP (lower) experiments across groups with or without *in vivo* binding sites (BS) and measured rG4 structures. Violin plots depict the data distribution, with box plots showing medians and interquartile ranges.

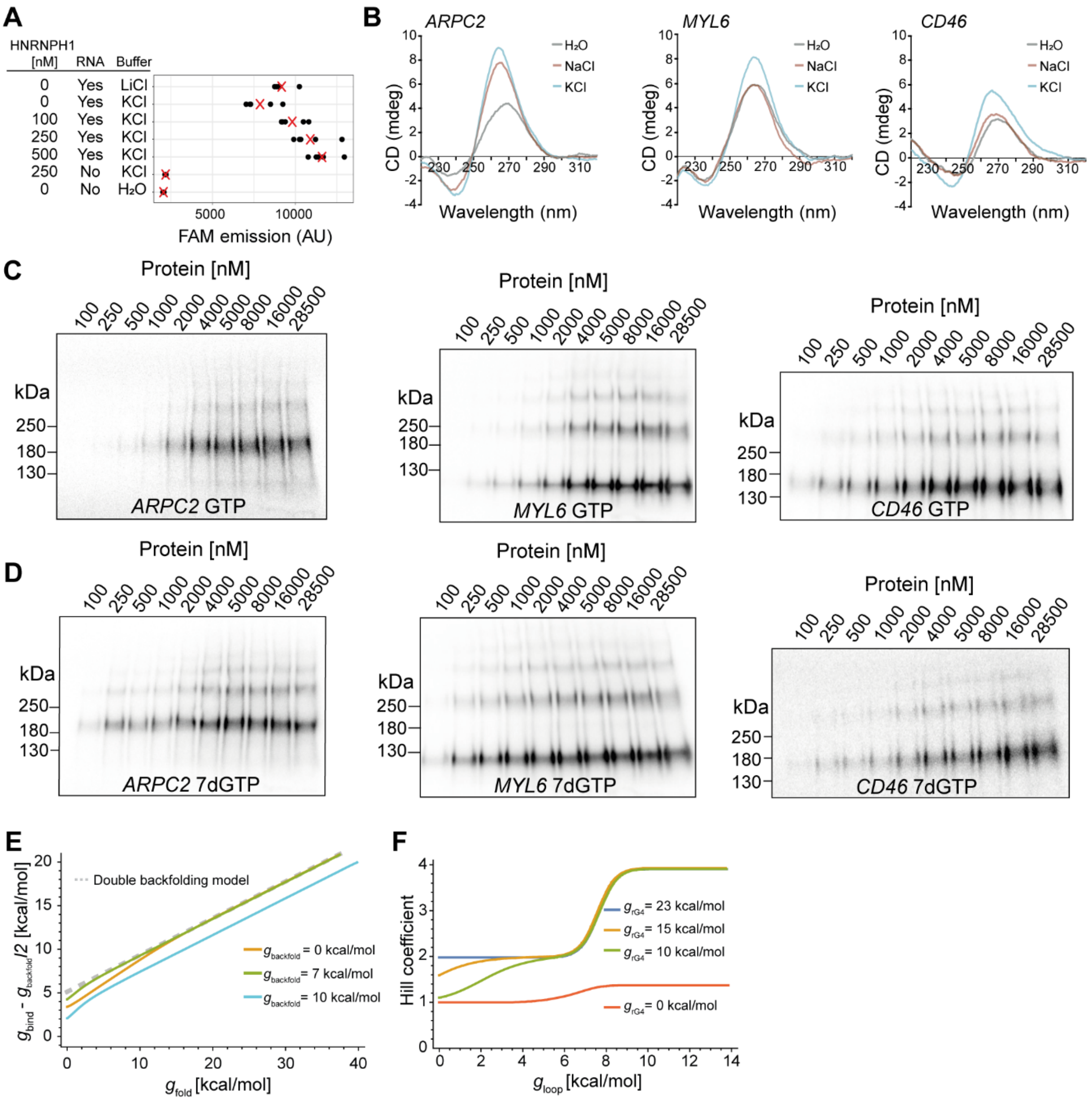

**Supplement Figure S5. Recombinant HNRNPH1 protein preferentially binds to rG4 RNAs in the unfolded state. (A)** Plot shows FAM emission of an rG4-forming FAM-TAMRA labeled RNA oligonucleotide upon addition of varying concentrations of recombinant HNRNPH1 protein, in the presence of 150 mM LiCl, 150 mM KCl, or no salt. FAM excitation was performed at 485 nm and FAM emission was measured at 520 nm. All experiments were performed in five individual replicates, represented by black dots, with red X representing mean. **(B)** Circular dichroism (CD) spectra for the RNA oligonucleotides *ARPC2*, *MYL6* and *CD46* (10 µM) at 25 °C to validate rG4 formation. CD spectra were measured in 20 mM Tris-HCl pH 7.5 buffer without salt ($H_20$) or in buffer with either 150 mM KCl or 150 mM NaCl. *ARPC2* and *MYL6* had a positive peak at ~264 nm and a negative peak at ~240 nm, while for *CD46* there was a peak shift to close to 270 nm for positive and ~245 nm for negative peak. **(C, D)** Autoradiograms of the *in vitro* titration series of recombinant HNRNPH1 protein with *CD46, MYL6,* and *ARPC2* RNA that was *in vitro* transcribed with GTP (C) or 7dGTP (D). **(E)** Reduced binding energy vs. folding energy needed to satisfy the constraint $K = 3 \cdot 10^{-6}$ mol/l. The dashed line shows the result for the double-backfolding model. **(F)** Hill coefficient vs. loop penalty for different folding energies and binding energy chosen such that $K = 3 \cdot 10^{-6}$ mol/l.

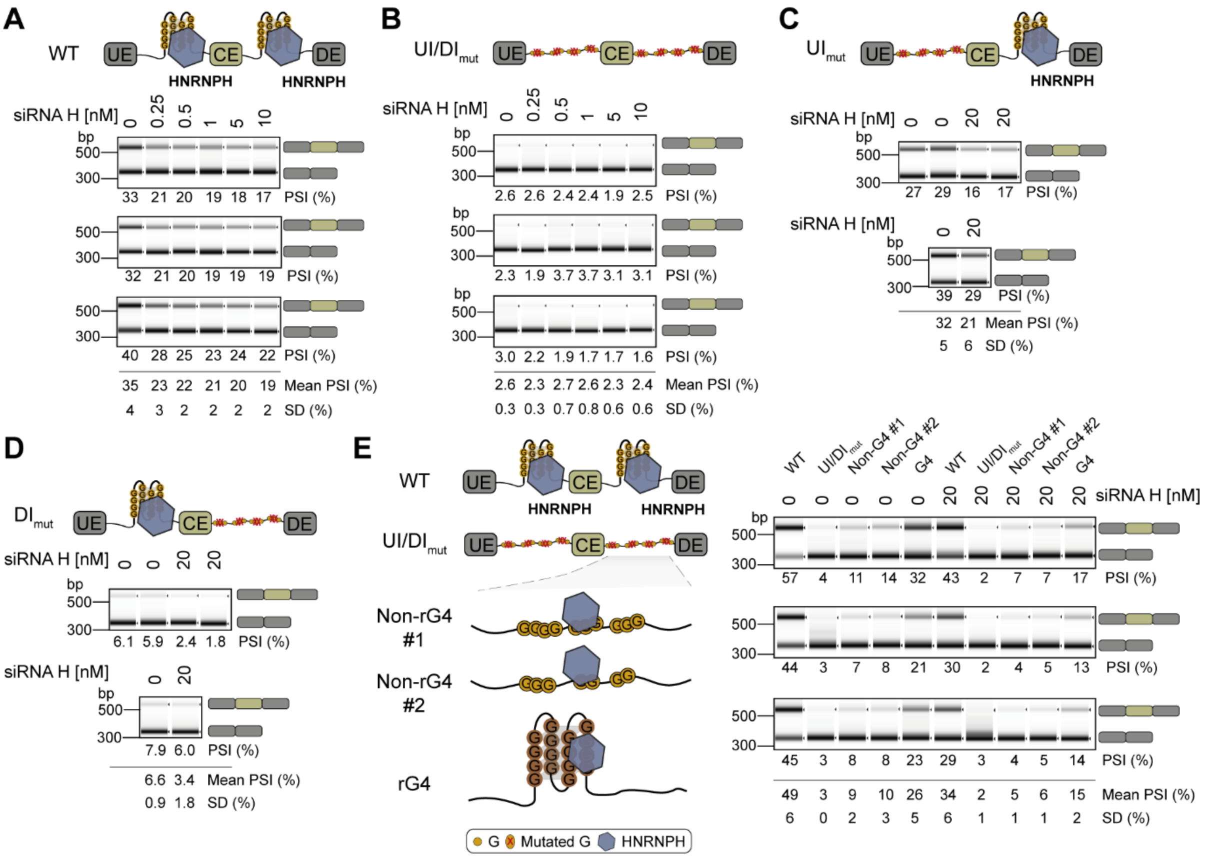

**Supplement Figure S6. AKR1A1 minigene splicing depends on HNRNPH binding sites. (A, B)** Semiquantitative RT-PCR is used to quantify the percent spliced-in (PSI) of *AKR1A1* exon 7 for the AKR1A1 WT (A), and UI/DI mutant (B) minigene in MCF7 cells upon gradual decrease of HNRNPH. The HNRNPH binding site locations in the genome are: chr1 45,568,349 to 45,568,431 (UI) and chr1 45,568,702 to 45,568,761 (DI), and were mutated by substitution of the middle G for each triple G-stretch (GGG), and the two middle Gs for each quadruple G-stretch (GGGG). Gradual reduction of HNRNPH was performed using increasing concentrations of a siRNA targeting *HNRNPH1* (siRNA H). Shown is the capillary gel electrophoresis of the RT-PCR products from three biological replicates with the respective PSI values below. Mean and standard deviation (SD) of the PSI across the replicates are depicted below. **(C, D)** Semiquantitative RT-PCR is used to quantify the percent spliced-in (PSI) of *AKR1A1* exon 7 for the AKR1A1 UI (C), and DI (D) mutant minigene in MCF7 cells upon *HNRNPH* KD (20 nM siRNA). Data representation as in (A, B). **(E)** Schematic illustration of the construct design for new HNRNPH binding sites in the downstream intron of the AKR1A1 minigene construct UI/DI. Two constructs with HNRNPH binding sites without rG4s (non-G4 #1 and #2) and one new construct with an HNRNPH binding overlapping with an rG4 (rG4) were generated (left). Semiquantitative RT-PCR is used to quantify the percent spliced-in (PSI) of *AKR1A1* exon 7 for the AKR1A1 for all illustrated mutants upon *HNRNPH* KD (20 nM siRNA). Data representation as in (A, B; right).

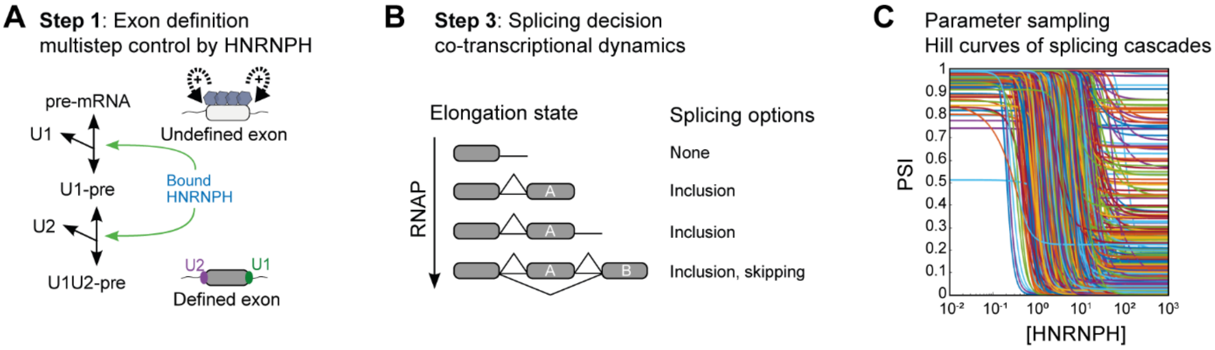

**Supplement Figure S7. Switch-like splicing regulation at multiple levels investigated by random parameter sampling. (A)** Switch-like control of exon definition by bound HNRNPH. Exon definition (step 2 in **Figure 6A**) occurs by stepwise binding of the pioneering U1 and U2 subunits of the spliceosome. If U1 and U2 binding is assumed to be cooperative and HNRNPH strongly enhances both binding steps, a Hill coefficient $n_H = 2$ is observed when plotting the defined state [U1U2-pre] as a function of the HNRNPH concentration (see Supplemental Material for details). **(B)** Switch-like exon inclusion arising from co-transcriptional splicing dynamics. The scheme shows the co-transcriptional nature of splicing decision-making (step 3 in **Figure 6A**): During transcript elongation by RNA polymerase II (RNAP), upstream sequences are available earlier when compared to downstream sequences, implying that splicing of the first intron (and thus a commitment for inclusion of the middle exon) is possible before skipping. Baeza-Centurion et al. (Baeza-Centurion et al. 2020) derived a kinetic model of cassette exon inclusion which describes how the PSI depends on the speed of transcript elongation and the relative strength (definition rates) of exons A and B (see **Supplemental Material**). When titrating the definition rate of exon A (% exon defined, x-axis) moderately switch-like behavior ($n_H = 1.4524$) can be observed in this co-transcriptional model (**Figure 6B** and **Supplemental Material**). **(C)** Analysis of switch-like HNRNPH-mediated splicing regulation for an *in silico* population of heterogeneous exons by parameter sampling. The splicing-regulatory cascade (**Figure 6A–C**) was repeatedly simulated while randomly sampling the kinetic parameters from log-normal distributions. Each line shows the sigmoidal dose-response curve for one parameter combination, and a corresponding histogram of Hill coefficients is shown in **Figure 6D**. See **Supplemental Material** for details.

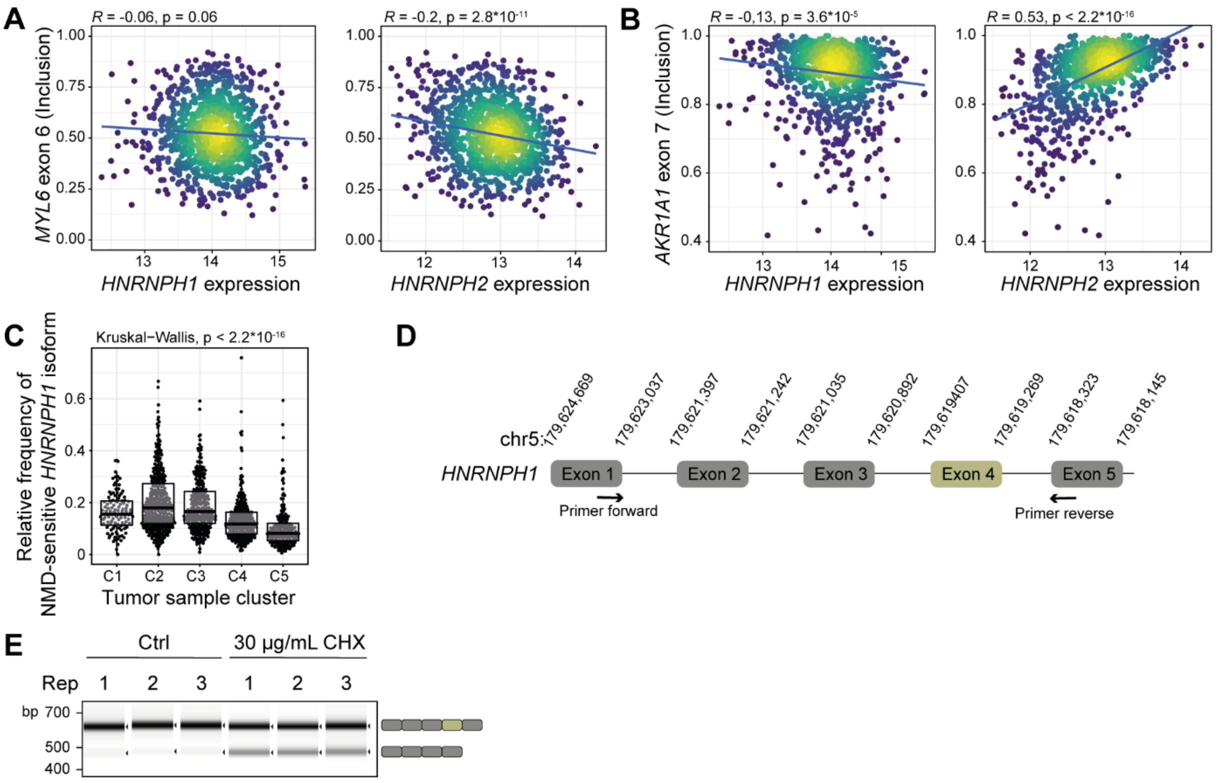

**Supplement Figure S8. HNRNPH-dependent splicing discriminates breast cancer subtypes, and its downregulation inhibits cell growth. (A, B)** Correlation of expression of *HNRNPH1* (left) and *HNRNPH2* (right) to inclusion of HNRNPH-regulated CE *MYL6* exon 6 (A) and *AKR1A1* exon 7 (B). Pearson correlation coefficient (*R*) and associated *P* value are shown. **(C)** Plot shows relative abundance of NMD-sensitive *HNRNPH1* exon 4 skipping isoform across five sample clusters (from *k*-means clustering in **Figure 7C**) representing tumor subtypes from C1 (WT) to C5 (most severe basal subtype). *P* value from Kruskal-Wallis test. **(D)** Schematic illustration of primer design strategy for confirmation of NMD-sensitive exon 4 skipping isoform of *HNRNPH1*. **(E)** Capillary gel electrophoresis of the RT-PCR products from three biological replicates in control (Ctrl) and NMD-repressing conditions (cycloheximide, CHX) for validating the NMD sensitivity of the *HNRNPH1* exon 4 skipping isoform using primers shown in (D).

### Supplementary Tables

**Table S1. Summary of high-throughput sequencing experiments.**

The table summarizes the number of reads for all high-throughput sequencing experiments conducted in this study. For RNA-seq (serial overexpression and knockdown of HNRNPH), iCLIP2, in vitro iCLIP2, and RTstop experiments, the individual replicates and their corresponding number of sequenced reads or read pairs are given. The number of uniquely mapped reads is additionally given for RNA-seq and iCLIP2 samples. For the iCLIP2 experiment, reads after duplicate removal are provided. Read pairs after adapter trimming are given for in vitro iCLIP2 and RTstop experiments.

**Table S2. Regulated alternative splicing events.**

The table lists all alternative splicing (AS) events detected across a titration series of HNRNPH KD and OE conditions. For each event, the table includes gene identifiers, genomic coordinates, and information about event type and complexity. PSI values (percent spliced in) are shown for each condition, along with corresponding ΔPSI values relative to control and posterior probabilities for differential splicing. Events are marked as reference-annotated or *de novo*. Each row is linked to a unique event_id and lsv_id for cross-referencing across datasets.

**Table S3. Fitted alternative splicing events.**

The table summarizes logistic fits for AS events that show a dose-responsive change across HNRNPH knockdown and overexpression conditions. For each event, the table reports the fitted minimum and maximum PSI values, the half-maximal effective concentration (EC50), the Hill coefficient ($n_H$), and the pseudo$R^2$ indicating fit quality. Events are classified into four categories based on their response dynamics: cooperative enhancement (coop-enh), cooperative repression (coop-rep), non-cooperative enhancement (non-coop-enh), and non-cooperative repression (non-coop-rep).

**Table S4. *In vitro* library construct information.**

The IVT library consists of 200-nt oligonucleotide sequences composed of an 18-nt T7 promoter, a 146-nt transcript region, a 15-nt unique barcode and a 21-nt L3 linker sequence for sequencing. The library covers 2,894 different transcript regions, including 1,340 HNRNPH binding sites with a predicted rG4 (BS+rG4), 1,409 HNRNPH binding sites without rG4 (BS-only) as well as 145 regions with predicted rG4s but no apparent HNRNPH binding (rG4-only). The 146-nt transcript regions were chosen such that the sites of interest (HNRNPH binding site or rG4) always start at position 51 in the 200-nt sequence. Each region is represented by four oligonucleotides with distinct barcodes, resulting in a total of 11,576 oligonucleotides. Barcodes were chosen to have a Hamming distance $\geq 5$ to any other barcode and contain no more than two consecutive Gs.

**Table S5. Results of RTstop and *in vitro* iCLIP2 experiments for each construct in the *in vitro* library.**

For each construct in the *in vitro* library, the following parameters are provided: RTstop classification (rG4/non-rG4), rG4 propensity, the number of G triplets, the presence of an HNRNPH binding site (true/false), rG4 prediction status, normalized *in vitro* iCLIP2 signals with GTP and 7dGTP, normalized RTstop signals with GTP and 7dGTP, and normalized HNRNPH binding strengths with GTP and 7dGTP.

**Table S6. List of oligonucleotides used in this study.**

Oligonucleotides used in this study include primer pairs for PCR amplification ("PCR"), oligonucleotides used for generating minigene reporter mutants ("Cloning"), oligos used for *in vitro* transcription ("*In vitro* transcription"), siRNAs ("siRNA"), and RNA oligos ("RNA oligos"). Name, sequence, and source are given for each oligonucleotide.

**Table S7. List of DNA plasmids used in this study.**

List of DNA plasmids used for series of minigenes experiments. For each plasmid, the corresponding plasmid name and the variation to the wildtype are described. The wildtype minigene reporter included *AKR1A1* exon 7 together with its flanking introns and exons. This alternative exon (AE), which is enhanced by HNRNPH (Dardenne et al. 2014; Zhang, Harvey, and Cheng 2019) displayed strong cooperativity in its dose-response and contained HNRNPH binding sites with overlapping rG4s in both flanking introns.

| No | Name | Backbone | Description |
|---|---|---|---|
| 1 | pcDNA3.1 (+)_EV | pcDNA3.1 (+) | Empty pcDNA3.1 (+) vector |
| 2 | HNRNPH1_OE | pcDNA3.1 (+) | Expression vector for *HNRNPH1* (Braun et al., 2018 (S. Braun et al. 2018)) |
| 3 | MBP-HNRNPH1-His | pET | *HNRNPH1* with N-terminal MBP-tag and C-terminal His-tag |
| 4 | AKR1A1 minigene | pcDNA3.1 (+) | *AKR1A1* minigene |
| 5 | AKR1A1 minigene_mutated 3'SS_2 | pcDNA3.1 (+) | *AKR1A1* minigene with G549T mutation |

| | | | |
|---|---|---|---|
| 6 | AKR1A1 minigene_mutated 3'SS_2_mutBS_UI | pcDNA3.1 (+) | *AKR1A1* minigene with mutated HNRNPH binding site in the upstream intron |
| 7 | AKR1A1 minigene_mutated 3'SS_2_mutBS_DI | pcDNA3.1 (+) | *AKR1A1* minigene with mutated HNRNPH binding site in the downstream intron |
| 8 | AKR1A1 minigene_mutated 3'SS_2_mutBS_UI-DI | pcDNA3.1 (+) | *AKR1A1* minigene with mutated HNRNPH binding site in upstream and downstream intron |
| 9 | AKR1A1 minigene_mutated 3'SS_2_noG4-BS-1_DI | pcDNA3.1 (+) | *AKR1A1* minigene with new HNRNPH binding site without rG4 in downstream intron |
| 10 | AKR1A1 minigene_mutated 3'SS_2_noG4-BS-2_DI | pcDNA3.1 (+) | *AKR1A1* minigene with new HNRNPH binding site without rG4 in downstream intron |
| 11 | AKR1A1 minigene_mutated 3'SS_2_G4-BS-1_DI | pcDNA3.1 (+) | *AKR1A1* minigene with new HNRNPH binding site with rG4 in downstream intron |
